# A Genetic Toggle Switch in Plants

**DOI:** 10.1101/2024.11.07.622546

**Authors:** Tessema K. Kassaw, Wenlong Xu, Christopher S. Zalewski, Katherine Kiwimagi, Ron Weiss, Mauricio S. Antunes, Ashok Prasad, June I. Medford

**Author notes:** These authors contributed equally to this work (Tessema K. Kassaw, Wenlong Xu). Sonata Therapeutics, Watertown, MA 02472. Front Range Biosciences, Boulder, CO 80026. Department of Biological Engineering, Massachusetts Institute of Technology, Cambridge, MA 02139. Department of Biological Sciences, University of North Texas, Denton, TX 76203. **Corresponding Authors:** June I. Medford, Department of Biology, Colorado State University, Fort Collins, CO 80523 USA; Ashok Prasad, Department of Chemical and Biological Engineering, Colorado State University, Fort Collins, CO 80523 USA.

## Abstract

In synthetic biology, genetic components are assembled to make transcriptional units, and transcriptional units are assembled into circuits to perform specific and predictable functions of a genetic device. Genetic devices have been described in bacteria, mammalian cell cultures and small organoids, yet development of programmable genetic circuits for devices in plants has lagged. Programmable genetic devices require defining the component’s quantitative functions. Because plants have long life spans, studies often use transient analysis to define quantitative functions while verification in stably engineered plants is often neglected and largely unknown. This raises a question if unique attributes of plants such as environmental sensitivity, developmental plasticity, or alternation of generations, adversely impacts predictability of plant genetic circuits and devices. Alternatively, it is also possible that genetic elements to produce predictable genetic devices for plants require rigorous characterization with detailed mathematical modeling. Here we use plant genetic elements with quantitatively characterized transfer functions and developed in silico models to guide their assembly into a genetic device: a toggle switch or a mutually inhibitory gene-regulatory device. Our approach allows computational selection of plant genetic components and iterative refinement of the circuit if the desired genetic functions are not initially achieved. We show that our computationally selected genetic circuit functions as predicted in stably engineered plants including through tissue and organ differentiation. Developing abilities to produce predictable and programmable plant genetic devices opens the prospect of predictably engineering plant’s unique abilities in sustainable human and environmental systems.

## INTRODUCTION

Plants naturally process information from biological and environmental inputs, using endogenous genetic networks that perform complex natural ‘computation’ through intricate, inter-twining genetic processes with multiple genes, input components, and genetic controllers. For example, plants receive input from diverse environmental sensors such as phytochromes (sensing red/far-red light) ^1, 2^ and cryptochromes (sensing blue light) ^3^. Plants produce Turing-like differentiation patterns that are computed with ligand-dependent controllers (e.g., auxin via TIR1; cytokinin via AHK4 receptors), and genetic On-Off switches that regulate plant states ^4–6^. Plant states include major developmental changes, embryonic and vegetative states, controlled by *WUSCHEL*, LEAFY allowing switching between vegetative and flowering states, and an epigenetic switch, FLOWERING LOCUS C, that regulates cold requirements for flowering ^7–9^.

Our understanding of natural switching of states in biology is based on experimental data combined with approaches that couple biological phenomena with quantitative analysis (e.g., *in vivo* measurement of gene expression and mathematical modeling). For example, the natural lysis-lysogeny switch in bacteriophages had been known for over a hundred years ^10^, but a detailed understanding required innovative mathematical models and experimental work that ultimately led to more refined mechanistic models ^11–13^. Likewise, work on the Lac operon from the early 1960’s proposed that genes are composed of genetic regulatory elements or “parts” capable of assembly into predictable genetic circuits ^14^. However, it was not until synthetic biology described the mathematical relationships between genetic elements that synthetic gene circuits could be developed and implemented ^15–17^. This practical theory allows both greater understanding of numerous gene regulation mechanisms and a new fundamental ability: to develop programmable genetic circuits with predictable function ^15, 16^. In microbes, such gene circuits can now be designed using automated software tools ^18^.

Predictable and programmable genetic circuits or “devices” for plants are important for developing environmentally sustainable technologies ^19–22^. Plant characteristics are unmatched by human manufacturing: plants have their own on-board energy system (photosynthesis), they self-assemble and self-repair and respond to environmental changes. Recent works on programming root development shows the challenges in developing plant technology and its potential power.

Brophy *et. al.*, developed a collection of synthetic transcriptional regulators for plants based on rapid prototyping using transient expression in tobacco leaves^23^. Using these genetic parts, they assembled genetic circuits to perform Boolean logic operations in plant roots and altered root architecture. Ragland *et. al.*, provide a recent review of synthetic biology efforts for engineering plant roots that includes addressing rhizobacterial interactions and biotic/abiotic stresses^24^.

In bacterial and mammalian systems, quantitative descriptions of genetic components allow construction of basic modules and circuits (*e.g.*, toggle switches, oscillators, sensors for cell-cell communication) ^15, 16, 25–27^. These modules can be combined and expanded to produce more sophisticated genetic devices (*e.g.*, delayed response systems, synchronized oscillators, systems to process signals and detect signal edges)^28–31^, production of designer metabolic networks, engineered T-cells and programmable organoids ^32–34^. If such technology were available for plants, we could design circuits for controllable induction of fruit ripening, detection systems for controlled activation of pathogen defense genes only when the pathogen is present, signal processing systems allowing plants to integrate endogenous and environmental signals for maximum yield, or designer networks for engineered production of plant bioproducts.

Forthcoming efforts could allow us to design complex fibers for specific applications (*e.g.*, cotton-plastic hybrids), engineered building materials formed from living plants that are capable of self-assembly and self-repair, novel systems for water treatment, along with major enhancement of the long promised renewable bioenergy sources through innovations such as self-shredding biomass.

A crucial need to achieve predictable and programmable genetic function is to quantitatively describe the behavior of a given genetic element (*e.g.*, promoter, activator, repressor) by characterizing an appropriate transfer function or input-output function. Gardner *et al.* first developed a synthetic genetic switch in *E. coli* by initially establishing the quantitative functions for promoters and ribosome binding sites (RBS) ^16^. These promoters/RBS/repressors were then rationally assembled into a genetic toggle switch consisting of two promoters and two repressors organized to control each other’s expression. The switch behavior results from the concentration of one repressor exceeding a threshold, and this accumulated repressor turns off expression of the other repressor. Importantly, genetic toggle switches can only exist in two stable states, and such bistable systems tolerate fluctuations associated with environmental conditions, gene expression or development of the organism.

We previously reported the construction and transfer function characterization of approximately 120 pairs of synthetic repressor-promoter for plants ^35^. Our components were characterized in protoplast assays with single-photon luciferase quantification and were found to function in both dicots (Arabidopsis) and monocots (Sorghum). Here, we detail how these components are used to rationally assemble a predictable and programmable genetic toggle switch in plants. Using our *in-silico* model, we selected two promoter-repressor pairs predicted to function as a toggle switch and assembled them into genetic circuits. *In planta* quantitative measurements showed that although these two circuits displayed some state switching behavior, neither achieved two stable and switchable states. Quantitative characterization of these two initial circuits allowed further refinement of our model, resulting in suggested circuit modifications that produced a functional toggle switch in stable plants. This work demonstrates that by integrating quantitative experiments and mathematical modeling, computer-selected genetic parts can be assembled into predictable and programmable regulatory circuits and devices in plants. In doing so, our work opens the possibility that in the future genetic circuitry design could be automated through machine learning.

## RESULTS

### Genetic toggle switch circuit architecture

The general design of our plant synthetic genetic circuit was initially based on the first genetic toggle switch demonstrated in bacteria ^16^. The ability to switch and maintain two distinct stable states is achieved with two constitutively active and repressible promoters that are mutually repressed by their corresponding repressor proteins (Figure 1A). However, bacterial systems, unlike plants, allow use of external inducers (e.g., IPTG) that directly bind to the opposite assembled and arranged in cistrons. To produce a switchable system in plants, we engineered repressors to be expressed as a result of transient application of chemical inducers, dexamethasone (DEX) or 4-hydroxytamoxifen (4-OHT) ^35^. Transcription of Repressor 1 is externally stimulated through a brief application of DEX, while transcription of Repressor 2 is externally stimulated through a brief application of 4-OHT (Figure 1A, Figure S1). Promoter 1 drives expression of Repressor 2, and is repressed by Repressor 1, whereas Promoter 2 controls expression of Repressor 1 and is repressed by Repressor 2. To obtain quantitative *in planta* measurement of our genetic circuit’s function, a second copy of Promoter 1 drives expression of the luciferase reporter (Figure 1A, Figure S1). Conventionally, toggle switches are defined as possessing two functional states: HIGH (ON) and LOW (OFF) ^16^. We use luciferase expression levels to define our two plant states: HIGH or ‘ON’ state and LOW or ‘OFF’ state ^16^ (Figure 1B & C). Fully operable toggle switches will maintain each state in the absence of the inducer, *i.e.*, show memory. Hence, switching states in our engineered plants should just need a brief chemical induction with the LOW or HIGH state maintained after induction (Figure 1A, Figure S1).

**Figure 1.**
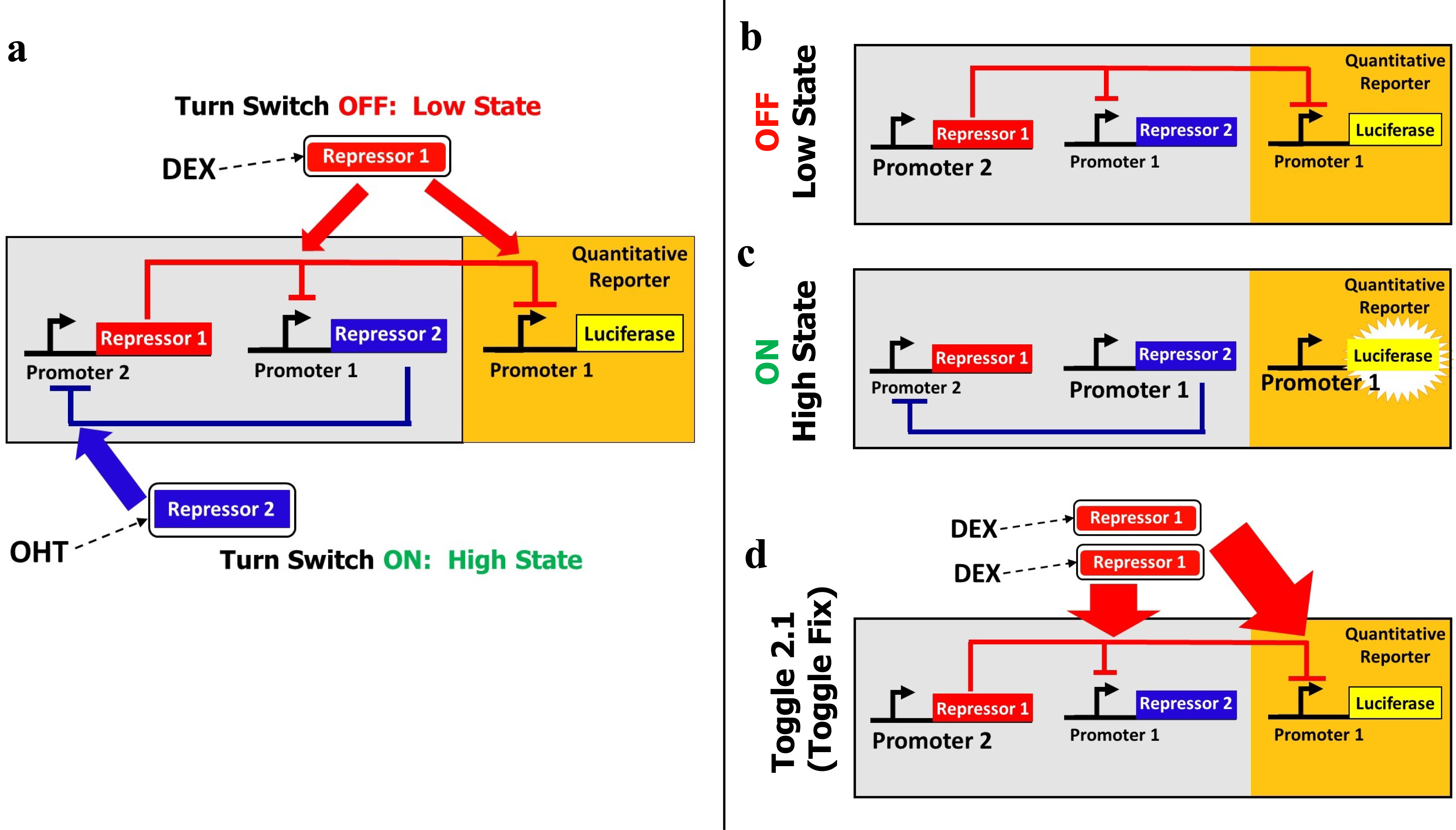
General architecture of a plant genetic toggle switch. (**a**) General architecture of the genetic toggle switch circuit. Repressor 1(red rectangle) inhibits transcription from Promoter 1. Repressor 2 (blue rectangle) inhibits transcription from Promoter 2. These two transcriptional units constitute the core of the toggle switch that enables maintenance of two states (large grey rectangle). A second copy of Promoter 1 drives the expression of a quantitative reporter, luciferase (yellow rectangle). The state of our plant circuit is described by the luciferase expression level: ON, High state, high expression; OFF, Low state, low expression. To switch the plant’s toggle state, externally activated ligand sensitive controllers to DEX or 4-OHT provide a transient chemical induction of Repressor 1 or Repressor 2, respectively (details in Figure S1). To switch the plant toggle state, we use two external controllers sensitive to either the DEX or 4-OHT ligand that provide a transient induction of Repressor 1 and Repressor 2, respectively (Fig. S1). In addition, DEX induces expression of the second copy of Repressor 1 that binds to and represses expression of Promoter 1 (red arrows) to turn the switch OFF. Likewise, OHT induces expression of a second copy of Repressor 2 that binds to and represses Promoter 2 (blue arrow) to turn the switch ON. **(b)** Plant toggle switch in the OFF or Low state. The toggle switch in the LOW state shows memory when, in the absence of the DEX ligand, continued expression of Repressor 1 inhibits expression of Promoter 1 (red lines), preventing luciferase expression. **(c)** Toggle switch in the HIGH state (showing memory). In the absence of the OHT inducer, continued expression of Repressor 2 inhibits expression of Promoter 2 (blue line), allowing continuous expression of luciferase. **(d)** Refinement of Toggle 2.0 to form the fully functional plant switch, Toggle 2.1, produced by the simple addition of one extra copy of DEX-inducible Repressor 1.

### State Switching

Plants with our genetic circuitry that are *briefly* exposed to DEX induce expression of the Repressor 1, which binds to and inhibits expression of Promoter 1 (red lines, Figure 1B, Figure S1), turning our engineered switch, and luciferase expression, OFF. Our genetic circuitry is designed to both enable switching to the LOW state and maintaining the LOW state without continuous or additional DEX induction (Figure S1). Conversely, when our engineered plants are briefly exposed to 4-OHT, the expression of Repressor 2 is induced, switching the circuit, and luciferase expression, ON. Repressor 2 binds to and inhibits Promoter 2 (blue lines, Figure 1C, Fig. S1). As with the LOW state, our genetic circuit is designed to enable a switch to the HIGH state and maintain the HIGH state without continuous or additional 4-OHT induction (Fig. S1).

### Genetic Component Selection

We designed our genetic components in a manner that allows us to ‘tune’ or modulate the function of promoters and repressors. We modulate activity of the repressible promoters by varying the number of DNA binding elements for the repressor, the spacing between DNA elements, and the locations of these relative to the transcription start site^35^. Because the backbone of repressible promoters consists of constitutively active promoters such as cauliflower mosaic virus (CaMV 35S), nopaline synthase (NOS) and figwort mosaic virus (FMV) ^36–38^, our library provides a means to quantitatively tune constitutively active and repressible promoters.

To engineer a library of synthetic repressors, we use translational fusions consisting of orthogonal DNA binding domains (LexA ^39^, Gal4 ^40^) with well characterized plant repressor domains. Plant transcriptional repressor domains used are the ethylene-responsive element binding factor–associated amphiphilic repression (EAR) domain, the B3 repression domain (BRD) ^41–44^, and a synthetic repressor domain, OFPx, consisting of a consensus sequence among repressor domains from natural OVATE (OFP) proteins ^35^. The highly modular design of our genetic components allows tuning of the genetic circuits by altering the number or composition of the constituent elements. For example, in Toggle 1, Promoter 2 is assembled with two copies of the LexA binding element. In contrast, in Toggle 2.0, Promoter 2 has four copies of the LexA binding element.

In our previous work, we characterized the transfer function of these modular genetic elements in plant cells at the single molecule level with luciferase using transient protoplast assays ^35^.

While this approach allowed us to avoid the long-time frames required for testing with transgenic plants, the quantitative parameters of the transfer functions derived from protoplast assays differed 0.2-2 -fold from those calculated when the same genetic elements were stably integrated in plants. Regardless of the source, this variance in our transient measurements will affect our ability to build genetic devices with predictable function (*i.e.*, using the data for predictive fits or *in silico* design). To account for this, we selected components based on a mathematical analysis of the conditions for bistability in intact plant systems (*SI Notes, Section 1*). Our analysis shows that a functional plant toggle switch requires repressor-promoter pairs with two properties: (i) the repressibility of a promoter by its cognate repressor should have high cooperativity (characterized by high Hill coefficients in the transfer function), and (ii) the two promoters must have high and balanced beta values which approximate maximal expression levels (Equation S1.

Having identified the needed characteristics, we selected components for assembly of a plant genetic toggle switch using transfer function data and these specific characteristics.

Significantly, by combining our mathematical analysis with our experimental quantitative data from plant cells, we were able to select the repressor-promoter pairs predicted to function as a plant genetic device, a bistable toggle switch ^35^.

### Production of Plant Genetic Devices

Based on criteria defined above, genetic components were assembled into two distinct toggle switch circuits, named Toggle 1.0 and Toggle 2.0. We initially chose a synthetic promoter that is assembled from the NOS Promoter with two copies of the Gal4 binding elements inserted downstream of the transcription start site, producing promoter pNOS_2XGal4_ (Figure S1-S2), simplified in Figure 1 to synthetic Promoter 1. Promoter 1 drives the expression of a synthetic transcriptional repressor, LEAR (Figure S1) or LOFPx (Figure S2) (translational fusions of LexA DNA binding domain to the EAR or OFPx repressor, respectively), simplified to synthetic Repressor 2 (Figure 1). Similarly, we assembled synthetic Promoter 2 using the backbone of the CaMV 35S promoter with two LexA-binding elements inserted downstream of the transcription start site (p35S_2XLexA_, Figure S1) for Toggle 1.0, or with four LexA-binding elements inserted in front of the TATA box (p35S_4xLexA_TATA, Figure S2) for Toggle 2.0. Promoter 2 drives the expression of synthetic Repressor 1, GEAR, assembled by linking the Gal4 DNA binding domain to the EAR repressor (Figure S1& S2). Promoter 1::Repressor 2 (blue rectangle) forms the first half or the first promoter-repressor pair of our toggle switch while Promoter 2::Repressor 1 (red rectangle) forms the second half or the second promoter-repressor pair (Figure 1). Collectively this produces two constitutively active and repressible synthetic promoters that are mutually repressed by each other’s cognate repressor. In both Toggle 1.0 and Toggle 2.0, a 2:1 combination of Promoter 2::Repressor 1 components was chosen to balance the maximal strength of the stronger Gal4-based promoter used in the Promoter 1 (see *SI Notes Section 2* and Table S10 for a detailed description of the quantitative characteristics of these components). We then evaluated the function of Toggle 1.0 (Figure S1) and Toggle 2.0 (Figure S2) in stably transformed plants.

### *In planta* Quantitative Measurement

In bacteria, data collection to assess genetic toggle circuit function benefits from short cell growth times (20-30 hours) ^16^ and high throughput methodology ^45^. For plants, determining circuit function requires non-destructive collection of quantitative data from the same plant over 10 or more days. Figure 2 describes our approach to non-destructively collect *in planta* luciferase expression data at the single molecule level while externally activating our genetic switch (see Materials and Methods). In brief, plants were grown vertically on MS agar plates for indicated times and transiently exposed to the external inducers to initiate the state switch. Single photon luminescence data are collected in a spatial pattern that corresponds to the plant (Figure 2A-B).

**Figure 2.**
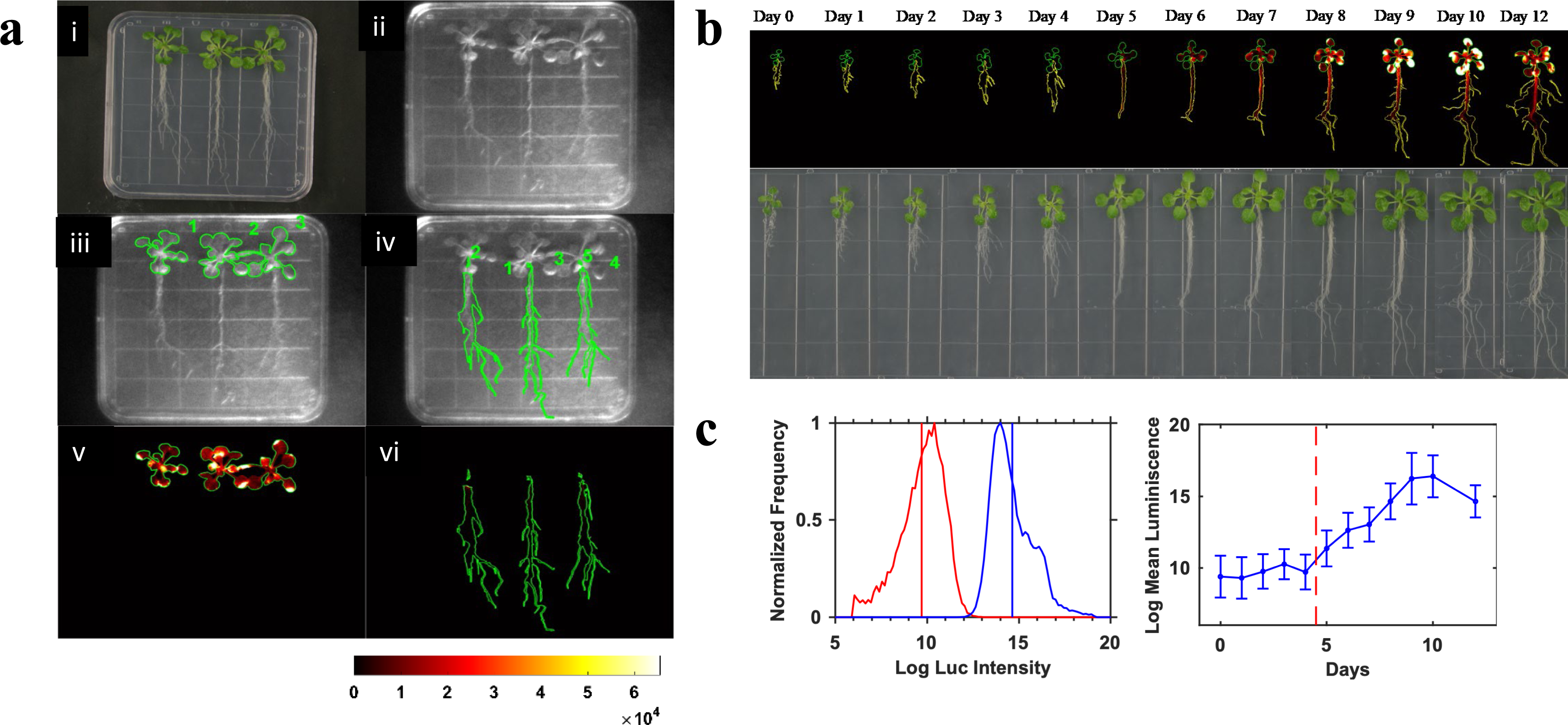
Non-destructive collection of single molecule data in whole plants with luciferase quantification. **(a)** Custom-designed program for whole plant image processing. A high-resolution color image is collected using a regular digital camera under well controlled lighting conditions prior to the luciferase imaging, which was used for color thresholding and identifying the regions of interest (ROIs) of shoots and roots for individual plants (*i*). Bright-field image collected *in situ* under the luciferase camera, which was used to align the ROIs obtained from *i* (*ii*). It should be noted that there are differences in resolution, orientation and lighting conditions between a and b. ROIs of shoots (*iii*) and roots (*iv)* aligned with the bright-field image with ROIs indexed spatially. ROIs of shoots (*v*) and roots (*vi*) applied to the luminescence images. **(b)** Representative luminescence images and pictures of Toggle 2.1 plants indicating the low to high switching. **(c)** Representative quantitative data of Toggle 2.1. The left panel shows histograms of pixel intensities of leaves at Day 0 (red) and at Day 12 (blue). These histograms are used to calculate the mean and standard deviation of the pixel luminescence. Right plot shows this for all 12 days.

Luminescence data was then superimposed with high resolution bright field images of the plants (Figure 2A), and a custom imaging processing computer script was developed (*Materials & Methods and SI Section 3*) that exacts the quantitative data from the spatially defined patterns.

Our custom program can distinguish data between shoots and roots (Figure 2A), allowing separate analysis of our genetic circuits in these plant organs.

### Initial States in Transgenic Plants

We used *in planta* luciferase expression measurements to obtain quantitative data describing the toggle circuit’s function over 12 days (Figure 2B & C). We first screened primary transgenic plants grown in the absence of any inducer and found plants in either the LOW or HIGH states (as determined by luciferase expression). We reasoned that the primary transgenic plants which are initially ON, in the HIGH state (strong luciferase expression), could either have a functional toggle with low dynamic range or they could simply have constitutive luciferase expression.

Hence, we selected plants that were initially OFF, in the LOW state (no luciferase expression), for further analyses. Transgenic plants that were OFF were allowed to self-fertilize and homozygous plants obtained for rigorous evaluation of our plant genetic circuits.

### State Determination in Plants

To determine whether our stably integrated genetic circuits function as a bistable toggle switch, we used an Ordinary Differential Equation (ODE) model of the genetic switch based on Hill functions to describe the input-output characteristics of the circuit (*See Materials and Methods*). We reasoned that we would be better able to parameterize the model if we demanded that the same set of parameters fit every experiment that we were able to carry out for a particular genetic circuit. We designed six such experiments that are described in the next section, each of which isolated a property of a genetic toggle switch. In order to fit the ODE model, we maximized a log-likelihood function that was defined as the negative logarithm of the sum-of-squared errors over the entire set of experimental conditions that the plant was subject to. Because we could have an infinite set of solutions (an under-determined system), we sampled the solution space using a random sampling strategy (Markov Chain Monte Carlo (MCMC) with the Metropolis algorithm^46^) to numerically maximize the likelihood function. We tested our method on simulated data and found that it accurately identified the true parameter values of the system (Figure 3). Parameter values that met the criteria of goodness of fit were then used to simulate the ODE model allowing us to assess the bistable properties of the system (*details Materials & Methods*).

**Figure 3.**
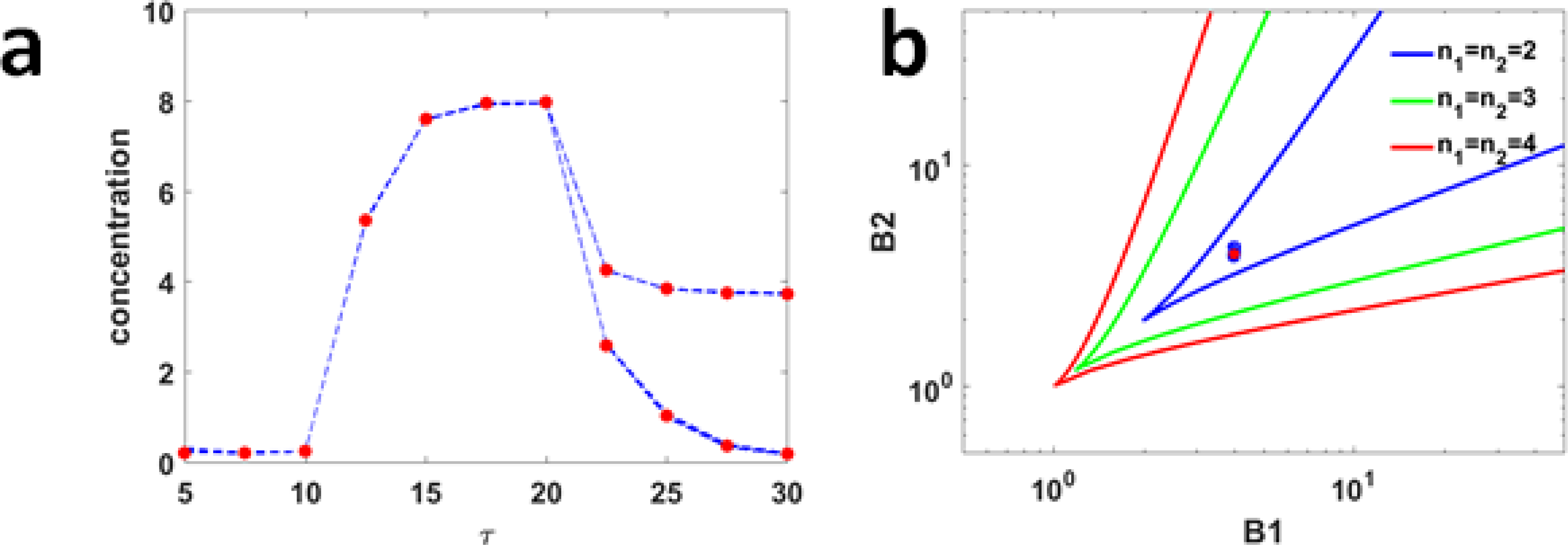
Reliability of our Mathematical approach as tested by simulated experimental data. (**a**) Simulated data and fitting the data to our model (MCMC fits). Numerical solutions of High memory and High to low as shown in red dots. MCMC fits are shown in blue dashed lines. (**b**) Simulated data and MCMC fits in 2D phase diagram of a plant toggle switch evaluated at different values of Hill coefficients (n). The axes b_x_ and b_y_ are dimensionless maximal expression levels of the repressible promoters (defined in **Equation S3)**. The bi-stable region is the region between the colored lines for each possible Hill coefficient. The Hill coefficients for both sides of the toggle were kept equal. True parameter value is shown by the red dot and the MCMC fits are the blue circles (n = 92).

### Testing the Plant Circuit

Prior to determining how our gene circuits function *in planta,* we first determined the time to establish steady-state behavior by quantifying the luciferase signal intensity after continuous induction with either inducer (DEX or 4-OHT). All genetic circuits, Toggle 1.0, Toggle 2.0 (and Toggle 2.1, below) established steady-state behavior in four days (Figure 4). Thus, this time point was used to define the time for plants to switch states (from the LOW State to the HIGH State and the HIGH State to the LOW State). We induced switching of circuit states after four days by moving plants to the opposite induction media (see *Materials and Methods*). We evaluated the genetic circuits for the following behaviors: [1] LOW Memory, *i.e.*, Maintenance of the genetic circuit in the OFF or LOW State, [2] Switch from LOW to HIGH State, or changing the genetic circuit from OFF to ON, [3] Switch from a HIGH to LOW State, or changing the genetic circuit from ON to OFF or [4] HIGH Memory, *i.e.*, Maintenance of the genetic circuit in the ON or HIGH State. In addition, each genetic toggle switch was evaluated under three additional conditions that served as Controls: no treatment Control, LOW State plants treated with inducer of the LOW State (control for OFF State), HIGH State plants treated with the inducer of the HIGH State (control for ON State). Representative images of whole plants under these experimental perturbations are shown in Figure S3 for Toggle 1.0.

**Figure 4.**
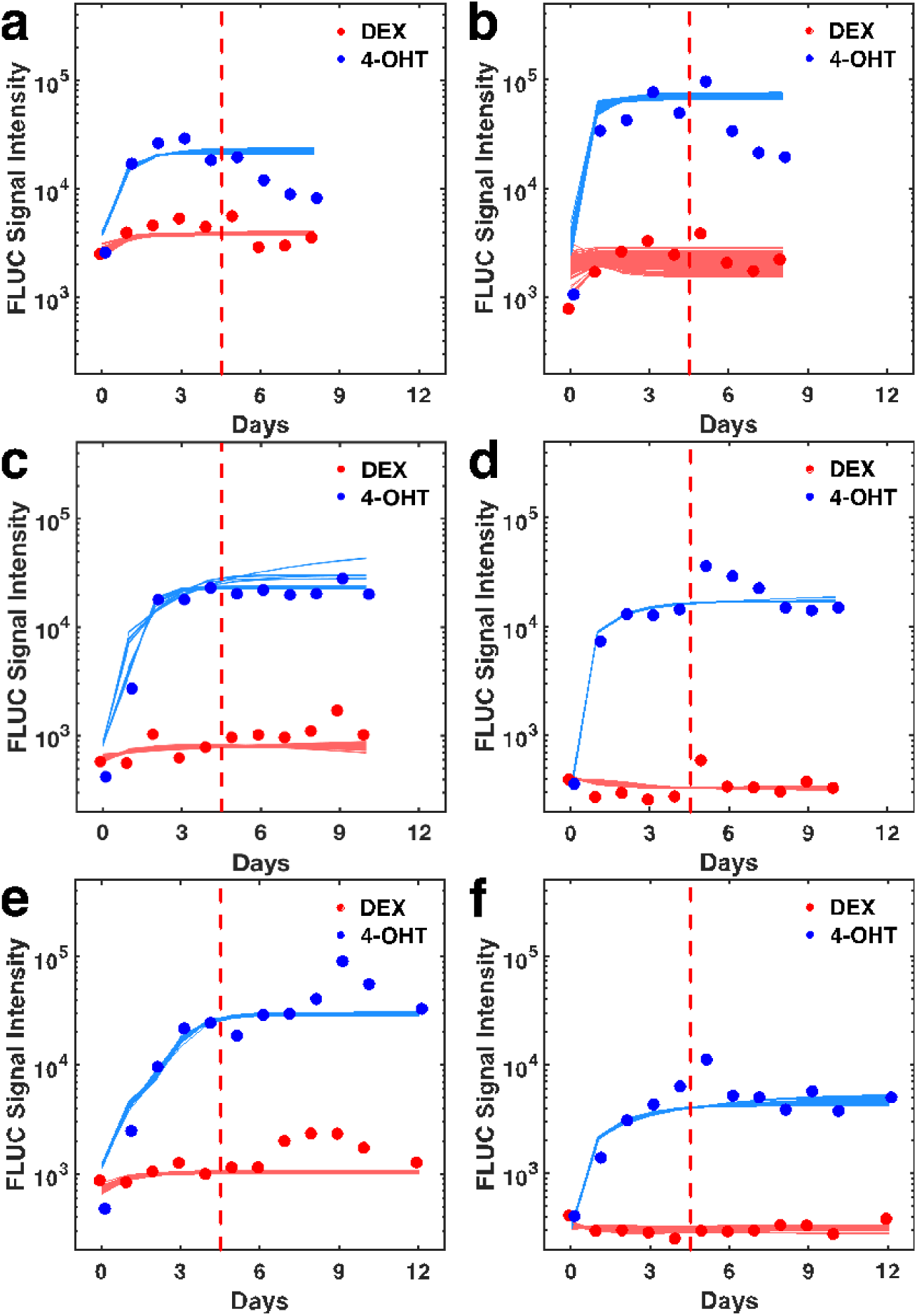
Steady state luciferase expression after continuous induction with 4-OHT or DEX. Shoots (**a**) and roots (**b**) of Toggle 1.0 steady state luciferase expression (*FLUC Signal Intensity*) over eight days of continuous induction. Shoots (**c**) and roots (**d**) of Toggle 2.0 steady state luciferase expression over ten days of continuous induction. Shoots (**e**) and roots (**f**) of Toggle 2.1 steady state luciferase expression over twelve days of continuous induction. Dots represent the experimental data and solid lines of the same color represent the model fitted to the experimental data. Each solid line represents a successful parameter set, as defined in the Materials and Methods. 4-OHT (blue dots/lines) switches the Toggle ON, to the HIGH state, whereas DEX (red dots/lines) switches the Toggle OFF, to the LOW state.

### Quantitative Characterization of Plants Containing Toggle 1.0 Circuit

A hallmark of a bistable toggle switch is memory, *i.e.*, the ability to maintain a stable state even in the absence of the inducer or other perturbations. Figure 5 shows luciferase measurement data and ODE model fitting of Toggle 1.0 and Toggle 2.0 circuits, using our MCMC method. Plants containing Toggle 1.0 are able to accurately switch from the LOW state (OFF) to the HIGH state (ON) (Figure 5A and Figure S4 & S5). Likewise, these plants are able to switch from the HIGH state (ON) to the LOW state (OFF) (Figure 5A); however, these plants are unable to show memory of the HIGH state (ON), *i.e.*, Toggle 1.0 plants are not able to keep luciferase expression ON in the absence of the inducer (Figure 5A, *HIGH MEMORY*). MCMC fitting analysis of Toggle 1.0 found that approx. 98% of all good parameter sets were monostable (SI Notes Section 3&4 and Figure S5). We therefore conclude that plants containing our Toggle 1.0 genetic switch lack bistability, and instead functioned as a simple inducible system similar to others previously described in plants ^47, 48^. Alternatively, this circuit’s behavior can be described as a monostable “doorbell” switch that stays in the ON state only while the inducer is present.

**Figure 5.**
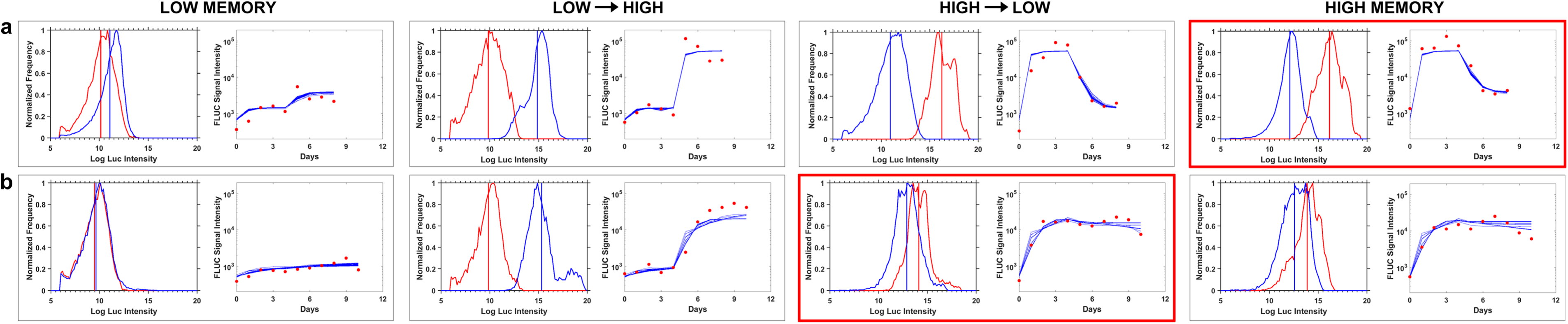
Quantitative analysis of bistability of toggle 1.0 and 2.0 in plants. Transgenic plants containing each of the toggle switch circuits were subjected to different inducer conditions (see Material and Methods for sample sizes) to test stability of the low and high states, as well as state transitions. Each box contains a histogram representation of the quantification of luciferase expression in plants, plotted as pixel intensity before (red curves) and after (blue curves) induction of state transition, and an ODE model fit (blue line) to the experimental data collected (red dots). **(*a*)** Toggle 1.0. **(*b*)** Toggle 2.0. Red boxes in ***a*** and ***b*** indicate where the corresponding toggle circuit failed to display the expected behavior. *Low Memory*, plants incubated with DEX inducer for 4 days to establish the low state, then moved to media with no inducer to test for stability (memory) of the low state. *Low → High*, plants incubated with DEX inducer for 4 days to establish the low state, then moved to media with 4-OHT inducer to switch the circuit to the high state. *High → Low*, plants incubated with 4-OHT inducer for 4 days to establish the high state, then moved to media with DEX inducer to switch the circuit to the low state. *High Memory*, plants incubated with 4-OHT inducer for 4 days to establish the high state, then moved to media with no inducer to test for stability (memory) of the high state.

### Toggle 2.0 Displays Bistable Function

In plants containing our genetic switch Toggle 2.0, Promoter 1.0 drives expression of the stronger LOFPx repressor (Repressor 2_OFPx_), and its cognate Promoter 2 has four LexA DNA binding sites inserted next to the TATA box region, Promoter 2_4Lex_ (Figure S2). Plants containing Toggle 2.0 are able to correctly switch from the LOW State (OFF) to the HIGH State (ON) with a brief exposure to 4-OHT, with an average induction of 37.5-fold in the shoots and 80-fold in roots (maximum 96-fold; Figure 5b, Figure S7). Further, unlike plants containing the Toggle 1.0 gene circuit, Toggle 2.0 shoots, but not roots, showed a memory ability (Figure 5b, Figure S7A & B and S8). However, Toggle 2.0 shoots (but not roots) poorly switched from the HIGH State (ON) to the LOW State (OFF) with DEX treatment (Figure 5B, right, Figure S7 and S8, and Tables S6 and S7). Our MCMC analysis found that 99.4% of the best parameter sets were bistable (*SI Notes Section 5*). Hence, in the shoots Toggle 2.0 has bistable properties, but appears unable to switch to the LOW State (OFF) with a brief exposure to DEX. Plants containing Toggle 2.0 show a pattern consistent with becoming trapped in the HIGH State (continuous luciferase production). Nonetheless, this type of genetic device could have plant biotechnology applications (see discussion).

Toggle 2.0 behavior was qualitatively different in the roots, and in fact despite a high fold change with induction of the HIGH state, it could not maintain this state (Figure S7 & S8). Toggle 2.0 could either be unable to switch because of an inducible promoter with insufficient strength or due to possible positional effects causing weak expression of the inducible promoter. We investigated the impact of positional effects from the T-DNA integration using thermal asymmetric interlaced polymerase chain reaction (TAIL-PCR) to retrieve sequences flanking insertion sites and estimate the number of insertions in plant chromosomes ^49^. We used TAIL-PCR to map the T-DNA insert to Chromosome 5 in the intergenic region between AT5G39980 and AT5G39990 (Figure S12 and Table S2-S3). As these genes are involved in chloroplast RNA processing and unidimensional cell growth, respectively, the chromatin surrounding Toggle 2.0 should not have unusual gene expression. This suggests that the genetic components of Toggle 2.0 are inadequate for a toggle switch function.

### Production of Fully Functional Toggle by Tuning Switching Dynamics

Because our approach used rigorous quantitative characterization of the genetic components and *in planta* function of the circuits, we were able to suggest modifications to the genetic circuit that are predicted to result in proper toggle function. Specifically, analysis of *in planta* function of Toggle 1.0 and Toggle 2.0 (Figure 5, SI *Notes*) suggests that we could produce a fully functional toggle switch by simply adding a second copy of the DEX-inducible GEAR repressor to Toggle 2.0 (Figure 1C, Figure S2).Therefore, we rapidly assembled this variant of the Toggle 2.0 genetic circuit, which we designated Toggle 2.1, indicating its minor iterative optimization (Figure 1D, Figure S2), and applied our experimental and mathematical analytical tools to determine its function in plants.

Figure 6 (and also Figure S9) shows quantitative data of luciferase expression in Toggle 2.1 plants over 12 days, whereas Figure 7 shows a mathematical analysis of the data (comparisons of experimental data with ODE simulations). Untreated plants containing Toggle 2.1 can maintain the LOW State throughout the 12-day course of our experiments (Figures 6A, 7C). In addition, Toggle 2.1 plants can properly switch from both states. Figure 6B (LOW→HIGH) and Figure 7C show Toggle 2.1 plants correctly switched from the LOW State to the HIGH State with 4-OHT addition, producing 34- and 19-fold increases in luciferase expression in the shoots and roots, respectively (Figure S10 C & D). Similarly, Figure 6C (HIGH→LOW) and Figure 7 show these plants are able to accomplish the HIGH→LOW switch. Lastly, Figure 6D and Figure 7 show that Toggle 2.1 plants are able to maintain the HIGH State (ON) throughout our 12-day experiment.

**Figure 6.**
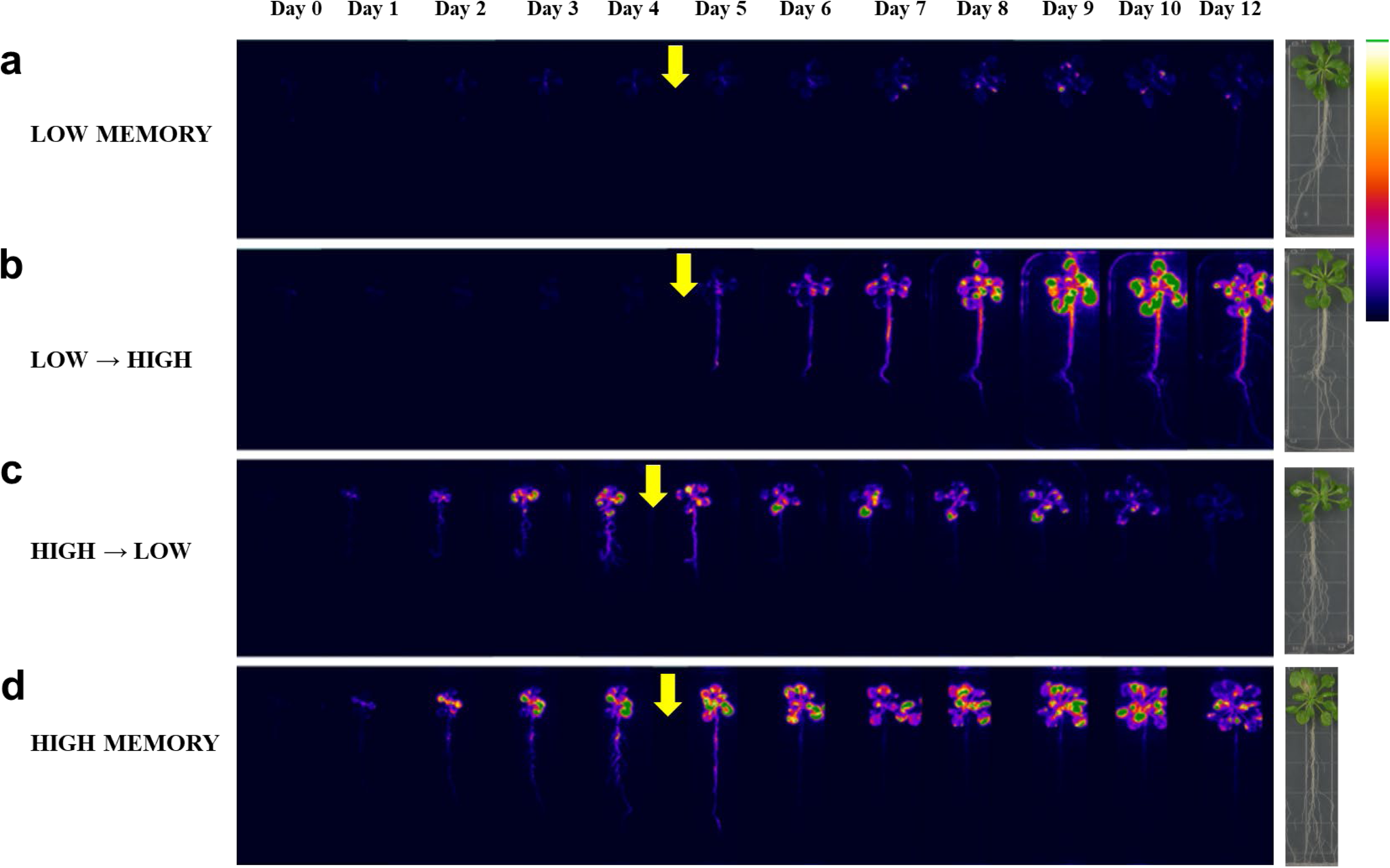
Single Photon Spatial Luciferase Data of plants containing Toggle 2.1. Homozygous transgenic plants containing the Toggle 2.1 circuit were grown vertically and imaged for luciferase over 12 days. The intensity of luciferase activity is false colored according to the fire scale (right). All plants were grown under conditions to establish steady state behavior for the plants genetic circuit. Yellow arrows indicate when plants were briefly exposed to conditions to switch the plant’s state. The HIGH state, induced with a brief exposure 4-OHT, switches luciferase expression ON. The LOW state, induced with a brief exposure to DEX, switches luciferase expression OFF. (**a) LOW Memory** retain OFF state through 12 days. After establishing the OFF steady state, plants are transferred to media without an inducer (DEX). The background or no luciferase expression indicates plants with Toggle circuit 2.1 have memory or (stability) of the LOW state. (**b) Switch Toggle to the HIGH State and remain ON.** After establishing the OFF steady state, plants are briefly exposed to 4-OHT to switch the gene circuit to the HIGH state. Plants keep luciferase expression ON through 12 days. (**c) Switch Toggle circuit to LOW and remain OFF.** After establishing the ON steady state, plants are briefly exposed to DEX to switch the gene circuit to the LOW state. Luciferase expression is rapidly turned OFF in roots and in shoots OFF through 12 days. (**d) High Memory** retain ON state through 12 days. After establishing the ON steady state, Plants are transferred to media with no inducer to test for stability (memory) of the high state. Plant maintain luciferase expression in shoots ON through 12 days.

**Figure 7.**
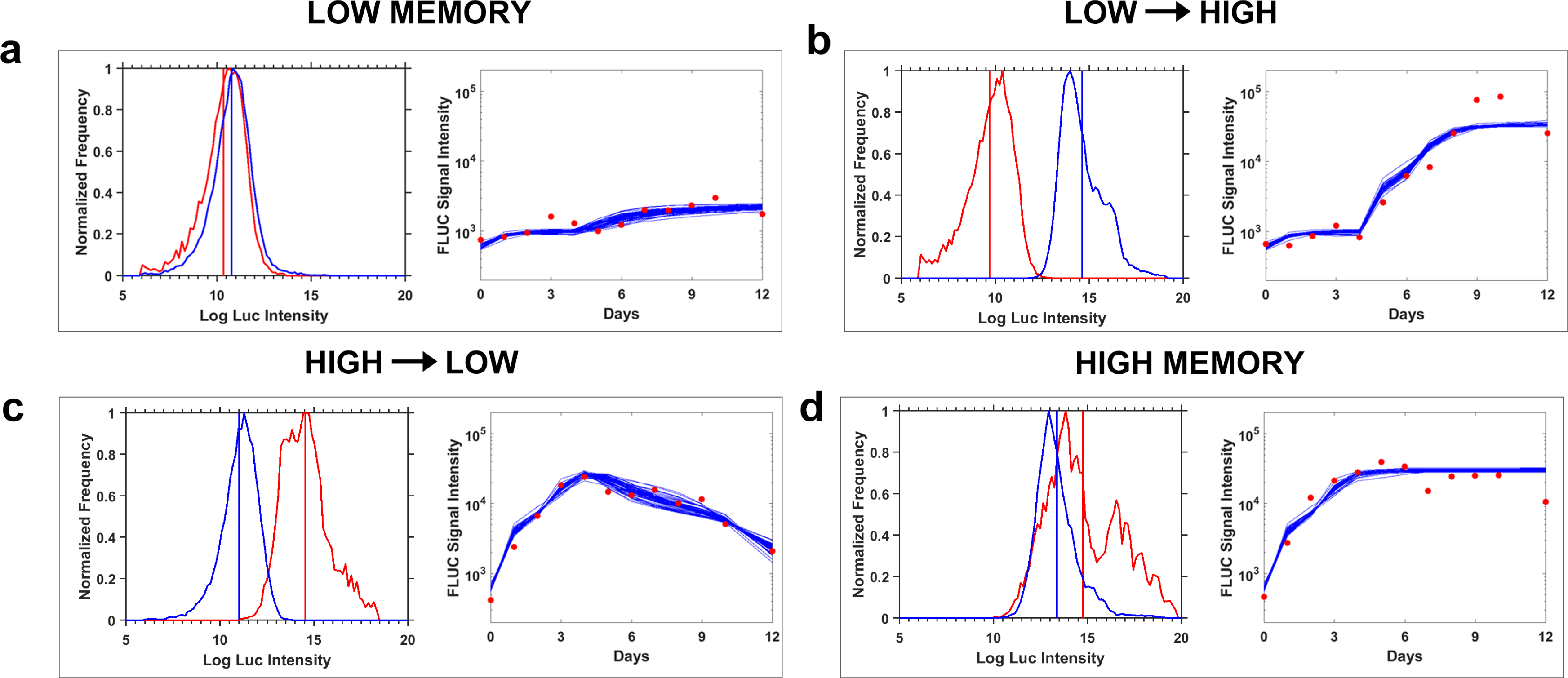
Toggle 2.1 is bistable. Transgenic plants containing Toggle 2.1 were subjected to different inducer conditions to test stability of the low and high states, as well as state transitions. Each box contains a histogram representation of the quantification of luciferase expression in plants, plotted as pixel intensity before (red curves) and after (blue curves) induction of state transition. Red points are data, blue lines are ODE model fits as described in *Materials and Methods*. (a) *Low Memory*, plants incubated with DEX inducer for 4 days to establish the low state, then moved to media with no inducer to test for stability (memory) of the low state. (b) *Low → High*, plants incubated with DEX inducer for 4 days to establish the low state, then moved to media with 4-OHT inducer to switch the circuit to the high state. (c) *High → Low*, plants incubated with 4-OHT inducer for 4 days to establish the high state, then moved to media with DEX inducer to switch the circuit to the low state. (d) *High Memory*, plants incubated with 4-OHT inducer for 4 days to establish the high state, then moved to media with no inducer to test for stability (memory) of the high state.

MCMC analysis was used to further quantitatively evaluate the bistability shown in Figure 6-7, showing that for shoots all the parameter sets that met the goodness of fit criterion are bistable, *i.e.*, they displayed two well-separated states for a time period much longer than the experimental timescale (*SI Notes Section 7*, and Figure S11). The MCMC analysis of the roots, while not as strong, still shows 53% to 32% of good parameter sets as long-lived well separated states (*SI Notes Section 8* and Fig. S12B). Plants containing Toggle 2.1 are able to switch between two distinct states and maintain these states, consistent with a fully functional toggle switch in plants.

## DISCUSSION

A fundamental aim of Synthetic Biology is to produce predictable and programmable traits in living organisms. Modular and predictable biological building blocks have been used for the construction of genetic circuits and biological devices that can be assembled to form complex information processing networks in microbes and mammalian systems ^15, 16, 25–29^. Our work developing a synthetic toggle switch in plants lays a foundation for both better understanding natural genetic circuits and producing predictable genetic control devices with considerable plant applications.

Previous studies have described various plant components (promoters, activators, repressors, etc.), as well as methods, principles and rules to rationally assemble them into genetic constructs ^50–52^. A recent plant study engineered a type of “memory” function using a bacteriophage recombination system ^53^. However, without quantitative analysis (transfer functions), which early synthetic biology research described as essential ^15, 16^, it is not possible to obtain predictable function nor use computational approaches for production of plant genetic circuits. We show that our quantitative predictions of component function, based on transient expression assays in protoplasts ^35^, can be adapted and used to guide the design of synthetic circuits in plants. In fact, our results show that the quantitative parameters that characterize genetic parts are robust across numerous experiments. Additionally, normalized parameter values estimated by the fits to the ODE model in whole plants overlap with those previously estimated for transient expression of the individual genetic parts in protoplasts (*SI Notes Sections 10* and Figure S13).

Our mathematical analysis also suggests methods to improve the ability of protoplast assays to predict function *in planta.* The analysis suggests that two uncharacterized factors in our approach, *noise* in the protoplast system (batch variability from protoplast prepared on different days) and *positional effects* from random integration of the T-DNA in whole plants (described here), may prevent complete predictability of our model. Future studies could use different transient expression methods or standardized integration of T-DNAs into specific chromosomal locations, or “landing pads”, typically regions of high transcriptional activity ^54–56^. It is notable that Toggle 2.0 integrated into a region not typically associated with high transcriptional activity (the modification to produce Toggle 2.1 was within the Toggle 2.0 genetic background).

When synthetic gene circuits are incorporated in multicellular organisms, an additional element of complexity is provided by differential gene expression in various tissues. In plants, one of the earliest differentiation events is distinction of the root from the shoot ^57^. When we quantitatively examined switching and memory in shoots and roots, we found differences in all our genetic toggle switches (Figure S5, S8, & S11). For instance, for Toggle 2.1, shoots appear to show both higher expression and better memory of the high state than roots (Figure 5; Figs. S10 and S11, and Tables S8 and S9). One possibility is that this arises because some of the viral promoters used as backbones for our synthetic circuit are known to have differential expression in roots ^37, 58, 59^. This could be addressed by separately doing quantitative analysis of genetic parts in protoplasts isolated from shoots and roots. A second possibility is that the absorptive properties of roots could disproportionally retain our inducers, making state switching/maintenance distinct from shoots. Another possibility is that positional effects from the random T-DNA integration ^60, 61^ may result in variable accessibility for transcription factors, potential for post-integration modifications, and differences in spatial and temporal regulation of promoter activity. In support of this, it is notable that the monostable toggle (Toggle 1.0) had higher expression in the roots than in the shoots (Figure S5). The distinct behavior of Toggle 1.0 versus Toggle 2.0 and Toggle 2.1 in roots versus shoots suggests that design of genetic circuits for multicellular and differentiated organisms could benefit by further quantitative tuning for specific tissues.

The importance of quantitative methods in synthetic biology has been previously emphasized ^15, 16^ and is illustrated here by our ability to identify the characteristics of each toggle switch design that we produced. The approach allowed us to diagnose the problem affecting Toggle 2.0 and suggest a minor iterative modification of the original design that resulted in full toggle function and bistability as predicted. Our measurements can also help identify what changes could be made for further improvement of Toggle 2.1, or to understand why Toggle 1.0 did not show bistability, if desired. For example, increasing the cooperativity of repression of Promoter 1 (pNOS_2xGal4_) by either adding extra transcriptional repressor binding elements (*e.g.*, 4xGal4) or changing the position of binding elements could further improve the switching dynamics of Toggle 2.1. Our work shows that quantitative experimental data coupled with mathematical modeling and model-aided design are powerful tools in developing programmable function of synthetic circuits in plants.

Toggle switches are both one of the first Synthetic Biology genetic devices developed ^16^ and a common component of electronics. The different toggle switches produced in our research could find application in plant biotechnology. For example, Toggle 2.0 retained the high expression state even without the inducer. This type of genetic device could be used to induce continuous synthesis of a specific product in field or other production settings in a controlled and regulated manner. Toggle 2.1, with its full bistable switching function, could be used to control a bottleneck in plant transformation, *i.e.*, switching to and from an embryogenic state ^62^.

Automated genetic design has already been demonstrated in bacteria ^18^ and it may be possible to do so with plants in the future. Our previous work showed that our components are functional in both monocots and dicots ^35^, hence producing predictable function with plants could be impactful to agriculture and a sustainable bioeconomy. Our toggle switch extends synthetic biology’s programmability to differentiated multicellular organisms. The genetic toggle switch maintains its function through meiosis, cell and tissue differentiation, and distinct life stages of plants, suggesting our synthetic circuits have robust genetic abilities. The functional retention through meiosis in plants, where the underlying gametophyte-sporophyte transition is accompanied by a dramatic change in chromatin accessibility and transcriptional reprogramming ^63^, is a significant milestone. The ability to produce programmable functions in plants may allow us to develop sustainable technologies for food security, materials, and devices to serve humanity and the environment.

## MATERIALS AND METHODS

### Plasmid Construction

All plasmids were constructed using standard molecular cloning techniques. Two types of plant transformation vectors were used for cloning (SI Appendix 1.1). The pCambia2300 binary vector (Kanamycin or BASTA resistant) was used to assemble the first toggle circuit, Toggle 1.0, in two separate T-DNAs. The pGREENII0229 vector was used to assemble Toggle 2.0 in one T-DNA. An additional DEX-inducible repressor to tune Toggle 2.0 was cloned in pCambia2300 and transformed into plants homozygous for Toggle 2.0 to create Toggle 2.1. The components used in individual toggle switch plasmids are described in SI Appendix, Fig. 1. To quantify the high/low switching behavior, the plant codon optimized luciferase gene from *Photinus pyralis* (firefly) was placed downstream of one copy of the repressible synthetic promoter, NOS_2xGal4_, in all three circuits. Primers and gBlocks were ordered from Integrated DNA technologies (IDT) and PCR reactions were performed with Herculase II (Agilent) or Phusion DNA polymerase (New England BioLabs) according to manufacturers’ protocols. In order to prevent transcription read-through among the genetic components, consecutive transcriptional units were insulated by transcription blocks^35, 64^. Plasmids were verified by DNA sequencing before using for plant transformation. DNA sequencing was provided by the Colorado State University Proteomics and Metabolomics Facility. Nucleotide sequences of individual parts used to assemble the toggle switch circuits are provided in SI Appendix, Table 1 and/or Schaumberg et al.^17^ The detailed design principles and quantitative characterizations of synthetic transcriptional repressor proteins and cognate repressible promoters were reported in Schaumberg et al. ^17^. For the schematic representation of the toggle switch circuits, see Fig. 2 and SI Appendix, Fig. 2.

### Plant Materials and Growth Conditions

*Arabidopsis thaliana* ecotype Columbia (Col-0) was used for all transformations. Plasmids were introduced into *Agrobacterium tumefaciens* GV3101 cells by electroporation. For Toggle 2.0 in the pGREENII0229 vector, the pSOUP helper plasmid was co-transformed. Stable transgenic plants were generated using the standard Agrobacterium floral dip method ^65^. Primary (T0) transgenic lines were selected on MS ^66^ media with 100 µg/ml Kanamycin, 34 µg/ml BASTA and 100 µg/ml cefotaxime (Toggle 1.0 and Toggle 2.1), or 34 µg/ml BASTA and 100 µg/ml cefotaxime (Toggle 2.0). Plants were screened initially for luciferase expression by spraying 500 µM d-luciferin solution in water (Gold Biotechnology, Inc.) containing 0.01% Tween 20, and imaging (described below). Homozygous plants were isolated at T3 (Toggle 1.0) or T2 generations (Toggle 2.0 and 2.1). For the luciferase assay, thirteen-day old plants from homozygous lines were transferred to MS media supplemented with 20 µM 4-OHT or 20 µM DEX for Toggle 1.0, and 5 µM 4-OHT or 50 µM DEX for Toggles 2.0 and 2.1, in 100 mm x 100 mm square plates (Fisher Brand, Cat No FB0875711A). Plants were grown vertically under short-day conditions (10 h light/14 h dark) at 22°C in a growth chamber. All experiments were conducted under seven different treatment conditions, including *control*, *Low → Low*, *Low Memory*, *Low → High*, *High → High*, and *High → Low*. ***Control***, plants incubated without inducer throughout the experiment. ***Low → Low***, plants incubated with DEX inducer throughout the experiment (low state). ***Low Memory***, plants incubated with DEX inducer for 4 days to establish the low state, then moved to media with no inducer to test for stability (memory) of the low state. ***Low → High***, plants incubated with DEX inducer for 4 days to establish the low state, then moved to media with 4-OHT inducer to switch the circuit to the high state. ***High → High***, plants incubated with 4-OHT throughout the experiment (high state). ***High → Low***, plants incubated with 4-OHT inducer for 4 days to establish the high state, then moved to media with DEX inducer to switch the circuit to the low state. ***High Memory***, plants incubated with 4-OHT inducer for 4 days to establish the high state, then moved to media with no inducer to test for stability (memory) of the high state. Both 4-OHT and DEX were obtained from Sigma-Aldrich.

### TAIL-PCR

Shoot and root genomic DNA was extracted from homozygous transgenic lines using the DNeasy plant mini kit (Qiagen) based on the manufacturer’s protocol. To identify the T-DNA insertion location in the Arabidopsis genome and the copy number of the transgene, thermal asymmetric interlaced polymerase chain reaction (TAIL-PCR) was employed ^26^. Three insertion-specific (nested) primers were designed to anneal to the left T-DNA border of the transgene and used in combination with arbitrary degenerate primers (AD primers) found in the literature (SI Appendix, Tables 2 and 3). Three subsequent TAIL-PCR reactions were performed and only the second- and third-round PCR products were analyzed on 1% agarose gels (SI Appendix, Fig. 4A). To recover unknown genomic sequences flanking the insertion, gel purified third-round TAIL-PCR products were either directly sequenced or cloned in pJET2.1 vector and sequenced using pJET-specific forward and reverse primers. Flanking sequences were identified by a BLAST search of the *Arabidopsis thaliana* Genome (TAIR10) to find the exact location of the transgene (SI Appendix, Fig. 4B). The transgene insertion was further confirmed by PCR amplification using insertion flanking primers.

### Image Collection and Processing

Luciferase activity was measured daily using single photon imaging with an XR/Mega-10 ICCD camera system (Stanford Photonics, Inc.) and available Piper software (v. 2.6.17) after spraying plants with 500 µM d-luciferin solution containing 0.01% Tween 20, and following a minimum of 30 minutes dark adaptation. Five data points (Days 0 to 4) were collected before switching inducers. Immediately before switching, plants were rinsed with MS liquid media for two hours by shaking. Four data points for Toggle 1.0 (Days 5 to 8), six for Toggle 2.0 (Days 5 to 10), or seven for Toggle 2.1 (Days 5 to 11) were collected after switching. To obtain plant boundaries for accurate quantification of the luciferase activity, a color high-resolution image was taken with a digital camera (Canon EOS Digital Rebel XTi) and a bright-field image was taken with the XR/Mega-10 ICCD camera immediately before measuring luciferase.

A custom designed image-processing software was developed to quantify the luminescence signals emitted from shoots and roots separately (Fig. S3). Briefly, the workflow consists of: 1) loading the high-resolution digital color image of the plants and indexing the regions of interest (ROIs) in shoots and roots for each plant through color thresholding (Fig. S3a); 2) carrying out an image registration step to align the color high-resolution and the bright-field images by user defined control points (n ≥ 2) (Fig. S3b). 3) masking the ROIs to the single photon measurements of luciferase activities, and making pixel-based measurements, including ROI areas in pixels, total intensity levels and a histogram of intensities (Fig. S3e-f); 4) exporting and logging measurement parameters and results for reproducibility. The image processing software is available on <u>GitHub.com</u> ^67^.

### Data Analysis and Fold Change Calculation

As we reported ^35^, the distribution of luciferase luminescence per tissue of an individual plant can be well described by a lognormal distribution (*i.e.*, a Gaussian distribution after a logarithmic transformation). The sample mean and standard deviations across plants in this study were therefore calculated on a logarithmic scale and then converted back to linear scale. Fold changes were calculated by dividing the mean luciferase data on the last day of the experiment by the one on the day immediately before changing an inducer. Differences in fold changes of two different treatments are statistically significant when the p-value is smaller than 0.05 by a 2-sample two-sided t-test.

### MCMC Parameter Fitting and Estimation of Bistability

Built on previously reported ordinary differential equation (ODE) models^16^ (*Equation 1*), our ODE model also consists of 1) two Hill functions describing the input-output characteristics of the two mutually repressible promoters; and 2) first-order dynamics for repressor degradation. Additional terms were introduced to better describe the behaviors of the toggle switches in plants, and address the increase in the complexity of the system: 1) binary terms (𝑘*_DEX_* and 𝑘*_𝑂HT_*) for the absence or presence of inducer; 2) constitutive expression terms (𝛼*_DEX_* and 𝛼*_𝑂HT_*) for the strengths of the inducible promoters; 3) 𝛼_𝑥_ and 𝛼_𝑦_ for the basal expression levels of the repressible promoters.

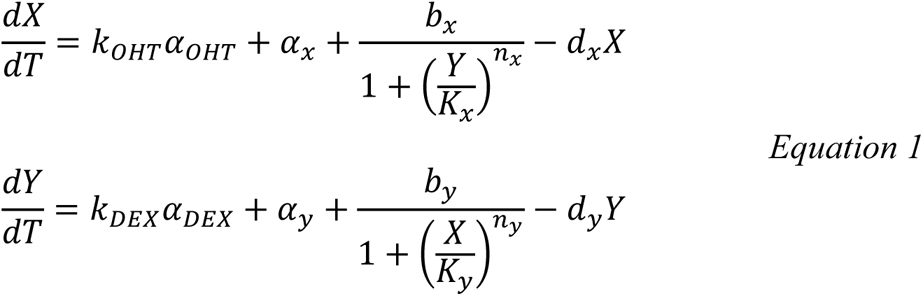

To test the bistability, we devised a novel method to fit the model (Equation 1) to the whole-plant luminescence data. We assumed the degradation constants of the two repressors are the same to simplify the fitting. We defined a likelihood function as the negative logarithm of the squared distances between the experimental data and numerical solutions of Equation S1, which is a weighted average of all seven different treatments in genetically identical plants with the same toggle circuit. We used a statistical approach, the Markov Chain Monte Carlo (MCMC) method, a class of algorithms for systematic random sampling of probabilities, to estimate the maximum-likelihood parameter values of the toggle switch tested in plants. Each MCMC run consists of: 1) Parameter initialization as a uniformly distributed random variable inside the parameter space (*e.g.*, Hill coefficients no greater than 6); 2) At each iteration, a Gaussian displacement was proposed for each parameter sequentially and the log likelihood function evaluated for every new set of parameter values. The proposed parameter set was accepted according to typical Metropolis criteria, where a proposed parameter value was either always accepted, if it increased the log likelihood, or accepted with a probability, if it decreased the log likelihood ^68^; 3) The MCMC run was terminated if either the maximum number of iterations (10,000) was reached, or a successful run with convergence achieved, defined as no significant change in log likelihood for more than 500 iterations. Only parameters obtained by a successful run were stored and retained for further analysis. The goodness of fit as measured by the value of the log likelihood function was bimodal in most cases, with a distinct population that clustered with the optimal fit. These were deemed the best parameter sets and were analyzed separately.

The fitting procedure was repeated at least 1,000 times (and up to 10,000 times) for the three toggle switch circuits to confidently sample the entire parameter space.

For the evaluation of bistability, the ODE model without the inducible terms (𝑘*_𝑂HT_*𝛼*_𝑂HT_* and 𝑘*_DEX_*𝛼*_DEX_*) was simulated numerically with the parameter sets. We initialized the model from two different conditions (high in one state and low in the other one), and the results were checked for bistability. We found that the numerical solutions fell in three classes: monostable, bistable or “indistinguishable from bistable”. Monostable systems reach the same steady state, and bistable ones result in two different steady state values. A parameter set that is “indistinguishable from bistable” produces two trajectories that remained separate and relatively stable (relative daily change smaller than 10%) for a time longer than the experimental timescale of 15 days. The bistable and “indistinguishable from bistable” parameter sets were together classified as “bistable”.

We carried out the fitting procedure to the data from both shoots and roots. We used the plant data in three different ways: (1) We fitted the mean of all the technical replicates of each experimental condition; (2) We fitted bootstrapped dataset with random selected technical replicate for each run and pooled the parameter sets; 3) We fitted the specific plants whose luminescence heatmap images are shown in Fig. S4, 7 &10.

The MCMC parameter fitting codes are available on GitHub.com ^67^.

## AUTHORS INFORMATION

### Author Contributions

J.I.M. and R.W. conceived the idea; M.S.A. developed the protoplast methodology; K.K. and A.P. did the initial data analysis and initial models; A.P. and W.X. developed advanced models; W.X. developed plant imaging automation software; M.S.A., T.K.K., C.S.Z. and J.I.M. designed the synthetic genetic circuits; T.K.K. and C.S.Z. assembled and verified all genetic circuits; T.K.K. and support staff produced all transgenic plants; and all authors aided in data analysis, preparation and writing of the manuscript.

### Notes

The authors declare competing financial interests: J.I.M. is the founder and President of Phytodetectors, and J.I.M., M.S.A., and T.K.K. are co-inventors on a patent application using plant genetic parts.

## ACKNOWLEDGMENTS

This study was supported by National Science Foundation grant CBET-0731029 to J.I.M. and R.W., by the U.S. Department of Defense, Defense Threat Reduction Agency (DTRA) grant W911NF-09-10526 to J.I.M., and by U.S. Department of Energy Advanced Research Projects Agency – Energy (ARPA-e) grant DE-AR0000311 to J.I.M., M.S.A. and A.P. We thank Dr. Diane McCarthy for help with preparation and editing of the manuscript and numerous undergraduates for help with plant growth and maintenance.

## REFERENCES

[1] Fankhauser, C., and Chory, J. (1997) Light control of plant development, Annu Rev Cell Dev Biol 13, 203–229.

[2] Rockwell, N. C., Su, Y. S., and Lagarias, J. C. (2006) Phytochrome structure and signaling mechanisms, Annu Rev Plant Biol 57, 837–858.

[3] Chapman, S., Faulkner, C., Kaiserli, E., Garcia-Mata, C., Savenkov, E. I., Roberts, A. G., Oparka, K. J., and Christie, J. M. (2008) The photoreversible fluorescent protein iLOV outperforms GFP as a reporter of plant virus infection, Proc Natl Acad Sci U S A 105, 20038–20043.

[4] Lau, S., Smet, I. D., Kolb, M., Meinhardt, H., and Jürgens, G. (2011) Auxin triggers a genetic switch, Nat. Cell Biol. 13, 611–615.

[5] Dharmasiri, N., Dharmasiri, S., Weijers, D., Lechner, E., Yamada, M., Hobbie, L., Ehrismann, J. S., Jürgens, G., and Estelle, M. (2005) Plant Development Is Regulated by a Family of Auxin Receptor F Box Proteins, Dev. Cell 9, 109–119.

[6] Pernisova, M., Grochova, M., Konecny, T., Plackova, L., Harustiakova, D., Kakimoto, T., Heisler, M. G., Novak, O., and Hejatko, J. (2018) Cytokinin signalling regulates organ identity via the AHK4 receptor in Arabidopsis, Development 145.

[7] Greb, T., Mylne, J. S., Crevillen, P., Geraldo, N., An, H., Gendall, A. R., and Dean, C. (2007) The PHD finger protein VRN5 functions in the epigenetic silencing of Arabidopsis FLC, Curr Biol 17, 73–78.

[8] Weigel, D., Alvarez, J., Smyth, D. R., Yanofsky, M. F., and Meyerowitz, E. M. (1992) LEAFY controls floral meristem identity in Arabidopsis, Cell 69, 843–859.

[9] Zuo, J., Niu, Q. W., Frugis, G., and Chua, N. H. (2002) The WUSCHEL gene promotes vegetative-to-embryonic transition in Arabidopsis, Plant J. 30, 349–359.

[10] Echols, H. (1986) Bacteriophage λ development: temporal switches and the choice of lysis or lysogeny, Trends in Genetics 2, 26–30.

[11] Arkin, A., Ross, J., and McAdams, H. H. (1998) Stochastic kinetic analysis of developmental pathway bifurcation in phage lambda-infected Escherichia coli cells, Genetics 149, 1633–1648.

[12] Weitz, J. S., Mileyko, Y., Joh, R. I., and Voit, E. O. (2008) Collective decision making in bacterial viruses, Biophys J 95, 2673–2680.

[13] St-Pierre, F., and Endy, D. (2008) Determination of cell fate selection during phage lambda infection, Proc Natl Acad Sci U S A 105, 20705–20710.

[14] Jacob, F., and Monod, J. (1961) On the regulation of gene activity, Cold Spring Harbor Symp. Quant. Biol. 26, 193–211.

[15] Elowitz, M. B., and Leibler, S. (2000) A synthetic oscillatory network of transcriptional regulators, Nature 403, 335–338.

[16] Gardner, T. S., Cantor, C. R., and Collins, J. J. (2000) Construction of a genetic toggle switch in Escherichia coli, Nature 403, 339–342.

[17] Cameron, D. E., Bashor, C. J., and Collins, J. J. (2014) A brief history of synthetic biology, Nat Rev Microbiol.

[18] Nielsen, A. A. K., Der, B. S., Shin, J., Vaidyanathan, P., Paralanov, V., Strychalski, E. A., Ross, D., Densmore, D., and Voigt, C. A. (2016) Genetic circuit design automation, Science 352, aac7341.

[19] McCarthy, D. M., and Medford, J. I. (2020) Quantitative and Predictive Genetic Parts for Plant Synthetic Biology, Frontiers in plant science 11.

[20] Medford, J. I., and McCarthy, D. M. (2017) Growing beyond: Designing plants to serve human and environmental interests, Current Opinion in Systems Biology 5, 82–85.

[21] Medford, J. I., and Prasad, A. (2016) Towards programmable plant genetic circuits, Plant J 87, 139–148.

[22] Medford, J. I., and Prasad, A. (2014) Plant synthetic biology takes root, Science 346, 162–163.

[23] Brophy, J. A. N., Magallon, K. J., Duan, L., Zhong, V., Ramachandran, P., Kniazev, K., and Dinneny, J. R. (2022) Synthetic genetic circuits as a means of reprogramming plant roots, Science 377, 747–751.

[24] Ragland, C. J., Shih, K. Y., and Dinneny, J. R. (2024) Choreographing root architecture and rhizosphere interactions through synthetic biology, Nature Communications 15, 1370.

[25] Kiani, S., Beal, J., Ebrahimkhani, M. R., Huh, J., Hall, R. N., Xie, Z., Li, Y., and Weiss, R. (2014) CRISPR transcriptional repression devices and layered circuits in mammalian cells, Nat Methods 11, 723–726.

[26] Slusarczyk, A. L., Lin, A., and Weiss, R. (2012) Foundations for the design and implementation of synthetic genetic circuits, Nature reviews. Genetics 13, 406–420.

[27] Basu, S., Gerchman, Y., Collins, C. H., Arnold, F. H., and Weiss, R. (2005) A synthetic multicellular system for programmed pattern formation, Nature 434, 1130–1134.

[28] Danino, T., Mondragon-Palomino, O., Tsimring, L., and Hasty, J. (2010) A synchronized quorum of genetic clocks, Nature 463, 326–330.

[29] Sohka, T., Heins, R. A., Phelan, R. M., Greisler, J. M., Townsend, C. A., and Ostermeier, M. (2009) An externally tunable bacterial band-pass filter, Proc Natl Acad Sci U S A 106, 10135–10140.

[30] Kramer, B. P., and Fussenegger, M. (2005) Hysteresis in a synthetic mammalian gene network, P Natl Acad Sci USA 102, 9517–9522.

[31] Davidsohn, N., Beal, J., Kiani, S., Adler, A., Yaman, F., Li, Y., Xie, Z., and Weiss, R. (2014) Accurate Predictions of Genetic Circuit Behavior from Part Characterization and Modular Composition, ACS Synth Biol.

[32] Xia, P.-F., Ling, H., Foo, J. L., and Chang, M. W. (2019) Synthetic genetic circuits for programmable biological functionalities, Biotechnol. Adv.

[33] Pols, T., Sikkema, H. R., Gaastra, B. F., Frallicciardi, J., Smigiel, W. M., Singh, S., and Poolman, B. (2019) A synthetic metabolic network for physicochemical homeostasis, Nat Commun 10, 4239.

[34] Guye, P., Ebrahimkhani, M. R., Kipniss, N., Velazquez, J. J., Schoenfeld, E., Kiani, S., Griffith, L. G., and Weiss, R. (2016) Genetically engineering self-organization of human pluripotent stem cells into a liver bud-like tissue using Gata6, Nat Commun 7, 10243.

[35] Schaumberg, K. A., Antunes, M. S., Kassaw, T. K., Xu, W., Zalewski, C. S., Medford, J. I., and Prasad, A. (2016) Quantitative characterization of genetic parts and circuits for plant synthetic biology, Nat. Methods 13, 94–100.

[36] Shaw, C., Carter, G., and Watson, M. (1984) A functional map of the nopaline synthase promoter, Nucleic acids research.

[37] Sanger, M., Daubert, S., and Goodman, R. (1990) Characteristics of a strong promoter from figwort mosaic virus: comparison with the analogous 35S promoter from cauliflower

[38] Benfey, P. N., Ren, L., and Chua, N. H. (1989) The CaMV 35S enhancer contains at least two domains which can confer different developmental and tissue-specific expression patterns, EMBO J. 8, 2195–2202.

[39] Schnarr, M., Oertel-Buchheit, P., Kazmaier, M., and Granger-Schnarr, M. (1991) DNA binding properties of the LexA repressor, Biochimie 73, 423–431.

[40] Giniger, E., Varnum, S. M., and Ptashne, M. (1985) Specific DNA binding of GAL4, a positive regulatory protein of yeast, Cell 40, 767–774.

[41] Wang, S., Chang, Y., Guo, J., Zeng, Q., Ellis, B. E., and Chen, J.-G. (2011) Arabidopsis Ovate Family Proteins, a novel transcriptional repressor family, control multiple aspects of plant growth and development, PLOS ONE 6, e23896.

[42] Wang, S., Chang, Y., Guo, J., and Chen, J.-G. (2007) Arabidopsis Ovate Family Protein 1 is a transcriptional repressor that suppresses cell elongation, Plant J. 50, 858–872.

[43] Tsukagoshi, H., Morikami, A., and Nakamura, K. (2007) Two B3 domain transcriptional repressors prevent sugar-inducible expression of seed maturation genes in Arabidopsis seedlings, Proc Natl Acad Sci U S A 104, 2543–2547.

[44] Ohta, M., Matsui, K., Hiratsu, K., and Shinshi…, H. (2001) Repression domains of class II ERF transcriptional repressors share an essential motif for active repression, The Plant Cell ….

[45] Barbier, I., Perez-Carrasco, R., and Schaerli, Y. (2020) Controlling spatiotemporal pattern formation in a concentration gradient with a synthetic toggle switch, Mol Syst Biol 16, e9361.

[46] Press, W. H., Teukolsky, S. A., Vetterling, W. T., and Flannery, B. P. (2007) Numerical Recipes: The Art of Scientific Computing, Cambridge University Press.

[47] Schena, M., Lloyd, A. M., and Davis, R. W. (1991) A steroid-inducible gene expression system for plant cells, Proc.Nat.Acad.Sci.USA 88, 10421–10425.

[48] Zuo, J., Niu, Q., and Chua, N. (2000) An estrogen receptor-based transactivator XVE mediates highly inducible gene expression in transgenic plants, Plant J.

[49] Singer, T., and Burke, E. (2003) High-throughput TAIL-PCR as a tool to identify DNA flanking insertions, Methods Mol Biol 236, 241–272.

[50] Belcher, M. S., Vuu, K. M., Zhou, A., Mansoori, N., Agosto Ramos, A., Thompson, M. G., Scheller, H. V., Loqué, D., and Shih, P. M. (2020) Design of orthogonal regulatory systems for modulating gene expression in plants, Nat. Chem. Biol. 16, 857–865.

[51] Vazquez-Vilar, M., Quijano-Rubio, A., Fernandez-del-Carmen, A., Sarrion-Perdigones, A., Ochoa-Fernandez, R., Ziarsolo, P., Blanca, J., Granell, A., and Orzaez, D. (2017) GB3.0: a platform for plant bio-design that connects functional DNA elements with associated biological data, Nucleic acids research 45, 2196–2209.

[52] Patron, N. J., Orzaez, D., Marillonnet, S., Warzecha, H., Matthewman, C., Youles, M., Raitskin, O., Leveau, A., Farré, G., Rogers, C., Smith, A., Hibberd, J., Webb, A. A. R., Locke, J., Schornack, S., Ajioka, J., Baulcombe, D. C., Zipfel, C., Kamoun, S., Jones, J. D. G., Kuhn, H., Robatzek, S., Van Esse, H. P., Sanders, D., Oldroyd, G., Martin, C., Field, R., O’Connor, S., Fox, S., Wulff, B., Miller, B., Breakspear, A., Radhakrishnan, G., Delaux, P.-M., Loqué, D., Granell, A., Tissier, A., Shih, P., Brutnell, T. P., Quick, W. P., Rischer, H., Fraser, P. D., Aharoni, A., Raines, C., South, P. F., Ané, J.-M., Hamberger, B. R., Langdale, J., Stougaard, J., Bouwmeester, H., Udvardi, M., Murray, J. A. H., Ntoukakis, V., Schäfer, P., Denby, K., Edwards, K. J., Osbourn, A., and Haseloff, J.(2015) Standards for plant synthetic biology: a common syntax for exchange of DNA parts, New Phytologist 208, 13–19.

[53] Bernabé-Orts, J. M., Quijano-Rubio, A., Vazquez-Vilar, M., Mancheño-Bonillo, J., Moles-Casas, V., Selma, S., Gianoglio, S., Granell, A., and Orzaez, D. (2020) A memory switch for plant synthetic biology based on the phage ϕC31 integration system, Nucleic Acids Res 48, 3379–3394.

[54] Danilo, B., Perrot, L., Botton, E., Nogue, F., and Mazier, M. (2018) The DFR locus: A smart landing pad for targeted transgene insertion in tomato, PLoS One 13, e0208395.

[55] Gaidukov, L., Wroblewska, L., Teague, B., Nelson, T., Zhang, X., Liu, Y., Jagtap, K., Mamo, S., Tseng, W. A., Lowe, A., Das, J., Bandara, K., Baijuraj, S., Summers, N. M., Lu, T. K., Zhang, L., and Weiss, R. (2018) A multi-landing pad DNA integration platform for mammalian cell engineering, Nucleic Acids Res 46, 4072–4086.

[56] Zhao, Y., Kim, J. Y., Karan, R., Jung, J. H., Pathak, B., Williamson, B., Kannan, B., Wang, D., Fan, C., Yu, W., Dong, S., Srivastava, V., and Altpeter, F. (2019) Generation of a selectable marker free, highly expressed single copy locus as landing pad for transgene stacking in sugarcane, Plant Mol Biol 100, 247–263.

[57] Goldberg, R. B., de Paiva, G., and Yadegari, R. (1994) Plant embryogenesis: zygote to seed, Science 266, 605–614.

[58] Amack, S. C., and Antunes, M. S. (2020) CaMV35S promoter – A plant biology and biotechnology workhorse in the era of synthetic biology, Current Plant Biology 24, 100179.

[59] Comai, L. (1990) Novel and useful properties of a chimeric plant promoter combining CaMV 35S and MAS elements, Plant molecular biology 15, 373.

[60] Matzke, A. J., and Matzke, M. A. (1998) Position effects and epigenetic silencing of plant transgenes, Curr Opin Plant Biol 1, 142–148.

[61] Gelvin, S. B. (2017) Integration of Agrobacterium T-DNA into the Plant Genome, Annu Rev Genet 51, 195–217.

[62] Altpeter, F., Springer, N. M., Bartley, L. E., Blechl, A. E., Brutnell, T. P., Citovsky, V., Conrad, L. J., Gelvin, S. B., Jackson, D. P., Kausch, A. P., Lemaux, P. G., Medford, J. I., Orozco-Cárdenas, M. L., Tricoli, D. M., Van Eck, J., Voytas, D. F., Walbot, V., Wang, K., Zhang, Z. J., and Stewart, C. N. (2016) Advancing crop transformation in the era of genome editing, The Plant Cell 28, 1510–1520.

[63] Borg, M., Papareddy, R. K., Dombey, R., Axelsson, E., Nodine, M. D., Twell, D., and Berger, F. (2021) Epigenetic reprogramming rewires transcription during the alternation of generations in Arabidopsis, eLife 10, e61894.

[64] Padidam, M., and Cao, Y. (2001) Elimination of Transcriptional Interference between Tandem Genes in Plant Cells, BioTechniques 31, 328–334.

[65] Clough, S. J., and Bent, A. F. (1998) Floral dip: a simplified method for Agrobacterium-mediated transformation of Arabidopsis thaliana, Plant J 16, 735–743.

[66] Murashige, T., and Skoog, F. (1962) A Revised Medium for Rapid Growth and Bio Assays with Tobacco Tissue Cultures, Physiologia Plantarum 15.

67. Prasad, A. (2024) https://github.com/prasadlabcsu/PlantToggleSwitch, Github.inc.

68. Press, W., Teukolsky, S., Vetterling, W., and Flannery, B. (2007) Numerical recipes: the art of scientific computing, 3rd Edition.

