## Supplementary material for "A Genetic Toggle Switch in Plants": S1

[1]These authors contributed equally to this work. [2]Corresponding authors. [3]Current address: Sonata Therapeutics, Watertown, MA 02472. [4]Current address: Front Range Biosciences, Boulder, CO 80026. [5]Current address: Department of Biological Engineering, Massachusetts Institute of Technology, Cambridge, MA 02139. [6]Current address: Department of Biological Sciences, University of North Texas, Denton, TX 76203.

**Corresponding Authors:**

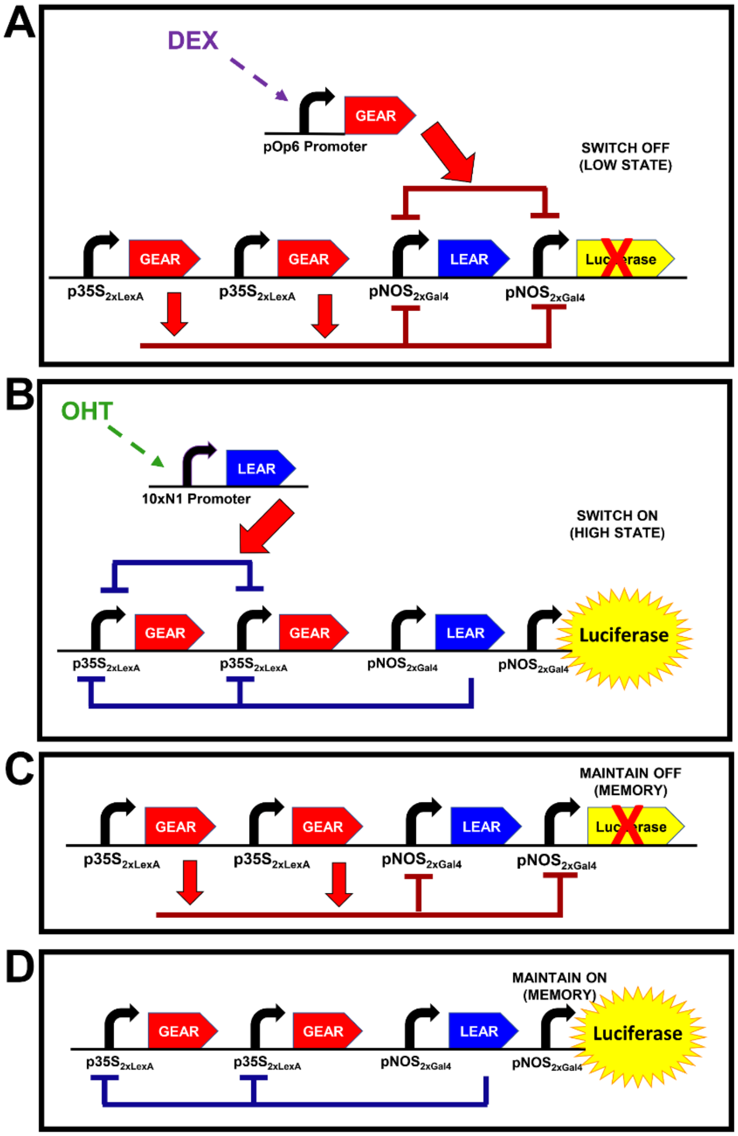


**Figure S1**. **Schematic showing details of Toggle 1.0 genetic components**. (**A**) External control of Repressor 1 transcription. DEX induces transcription of the pOp6 promoter activating expression of a copy of the repressor, GEAR, which is in addition to the main toggle components shown in Figure 1. The synthetic repressor, (GEAR) is composed of Gal4 DNA binding sites and the EAR repressor domain. GEAR binds to the constitutively active and repressible promoter $NOS_{2xGal4}$, or Promoter 1 in Figure 1, the main text. Promoter 1, $pNOS_{2xGal4}$, drives expression of both luciferase and Repressor 2. Hence, a brief exposure to DEX is designed to switch the genetic circuit to the LOW State or OFF. (**B**) External control of Repressor 2 transcription. OHT induces transcription of the 10xN1 promoter activating expression of a copy of the repressor, LEAR, which is in addition to the main toggle components shown in Figure 1. The synthetic repressor, Repressor 2 (LEAR) is composed of LexA DNA binding sites and the EAR repressor domain. LEAR binds to the constitutively active and repressible promoter, $p35S_{2xLexA}$, or Promoter 2 in the main text, Figure 1. Hence, a brief exposure to OHT is designed to switch the genetic circuit to the HIGH State or ON. (**C**) Details of genetic components designed to maintain the Toggle in the LOW state or off (corresponding to main text Figure 1B). In the absence of

DEX, Promoter 2 ($p35S_{2xLexA}$) continues to direct production of the GEAR repressor keeping expression of the LEAR repressor and luciferase OFF. (**D**) Details of the genetic components maintaining the Toggle in the HIGH state or on (corresponding to main text Figure 1C). In the absence of OHT, Promoter 1 ($NOS_{2xGal4}$) continues to direct expression of the LEAR repressor keeping expression of the GEAR repressor OFF while allowing luciferase expression.

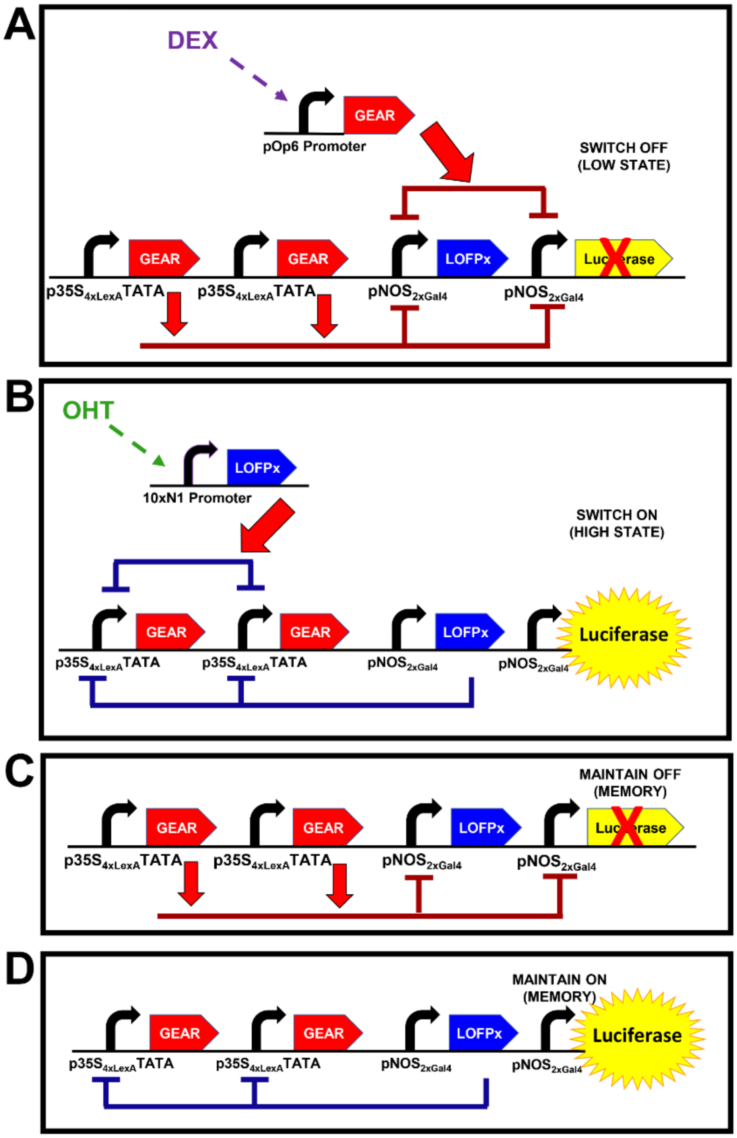


**Figure S2**. **Schematic showing details of Toggle 2.0 genetic components**. (**A**) External control of Repressor 1 transcription. DEX induces transcription of the pOp6 promoter activating expression of a copy of the repressor, GEAR, which is in addition to the main toggle components shown in Figure 1. As in Toggle 1.0, the synthetic repressor, (GEAR) is composed of Gal4 DNA binding sites and the EAR repressor domain. As in Toggle 1.0, GEAR binds to the constitutively active and repressible promoter $NOS_{2xGal4}$, or Promoter 1 in Figure 1, the main text. Also, like Toggle 1.0, Promoter 1, $pNOS_{2xGal4}$, drives expression of both luciferase and Repressor 2. Hence, a brief exposure to DEX switches the genetic circuit to the LOW State or OFF. (**B**) External control of Repressor 2 transcription. OHT induces transcription of the 10xN1 promoter activating expression of a copy of the repressor, LOFPx, which is in addition to the main toggle components shown in Figure 1. The synthetic repressor, Repressor 2 (LOFPx) is composed of LexA DNA binding sites and the OFPx repressor domain. LOFPx binds to the constitutively active and repressible promoter, $35S_{4xLexA}$TATA, or Promoter 2 in the main text, Figure 1. Hence, a brief exposure to OHT switches the genetic circuit to the HIGH State or ON. (**C**) Details of genetic components designed to maintain the Toggle in the LOW state or off (corresponding to main text Figure 1B). In the absence of DEX, Promoter 2 continues to direct production of the GEAR repressor keeping expression of the LOFPx repressor and luciferase OFF. (**D**) Details of the genetic components maintaining the Toggle in the HIGH state or on (corresponding to main text Figure 1C). In the absence of OHT, promoter 1 continues to direct expression of the LOFPx repressor keeping expression of the GEAR repressor OFF while allowing luciferase expression. Toggle 2.1 contains a second copy of the external controller, the DEX inducible GEAR repressor, Repressor 1.

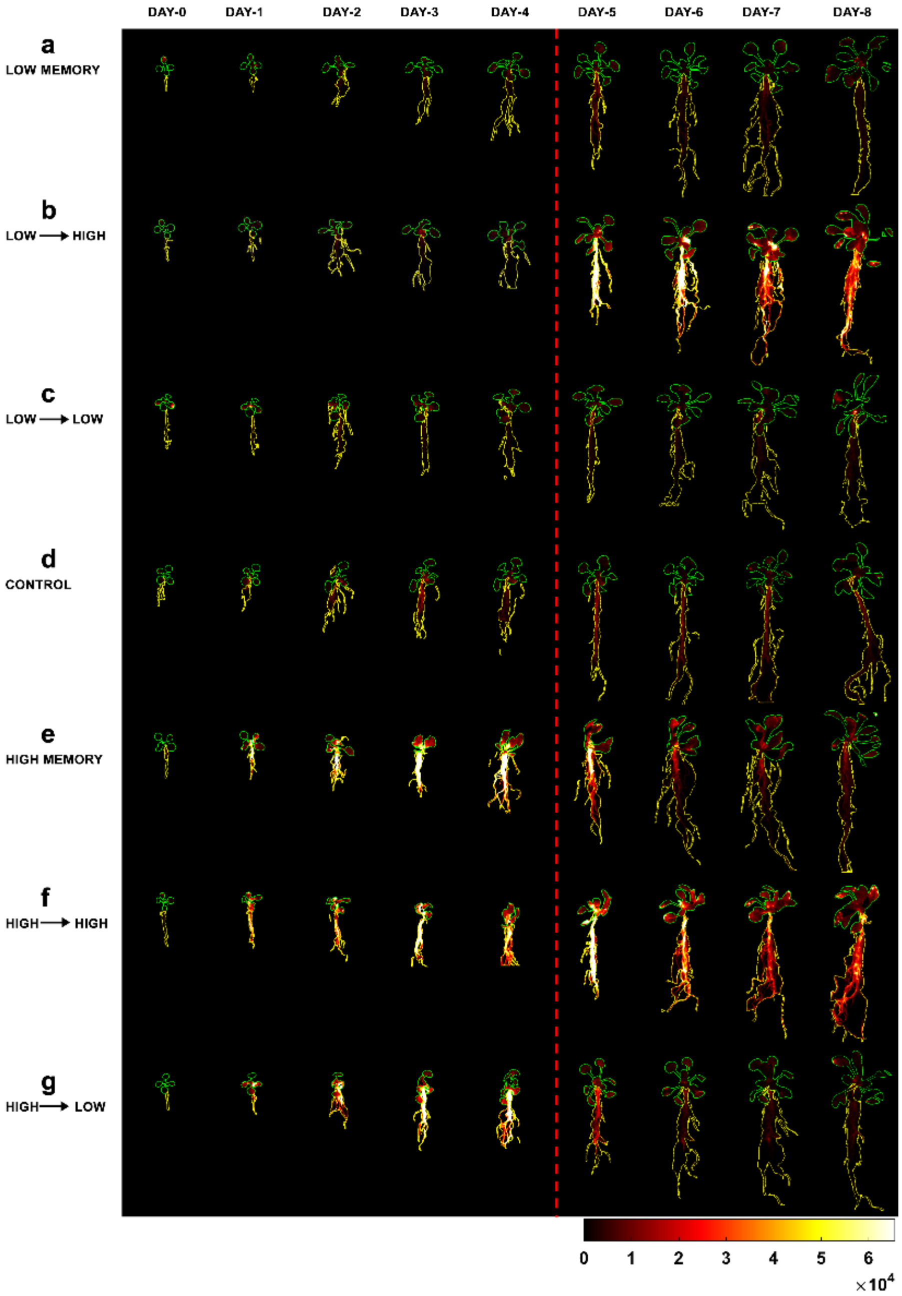


**Figure S3. Luminescence heat maps (linear scale) of selected plants containing Toggle 1.0 under different treatment conditions**. Regions of Interest (ROIs) drawn for the shoots are indicated by green lines, whereas those for the roots are indicated by yellow lines. All images use the same fire scale which false-colors luminescence intensity, with black representing zero and white representing saturation according to the scale shown at the bottom. The colors reflect luminescence intensity changes between different treatments. Inducer conditions were changed after imaging on Day 4, which is indicated by the red dashed line. **(a)** ***Low Memory***, plants incubated with DEX inducer for 4 days to establish the low state, then moved to media with no inducer to test for stability (memory) of the low state. (**b**) ***Low → High***, plants incubated with

DEX inducer for 4 days to establish the low state, then moved to media with 4-OHT inducer to switch the circuit to the high state. (**c**) ***Low → Low***, plants incubated with DEX inducer throughout the experiment (low state). (**d**) ***Control***, plants incubated without inducer throughout the experiment. (**e**) ***High Memory***, plants incubated with 4-OHT inducer for 4 days to establish the high state, then moved to media with no inducer to test for stability (memory) of the high state. (**f**) ***High → High***, plants incubated with 4-OHT throughout the experiment (high state). (**g**) ***High → Low***, plants incubated with 4-OHT inducer for 4 days to establish the high state, then moved to media with DEX inducer to switch the circuit to the low state.

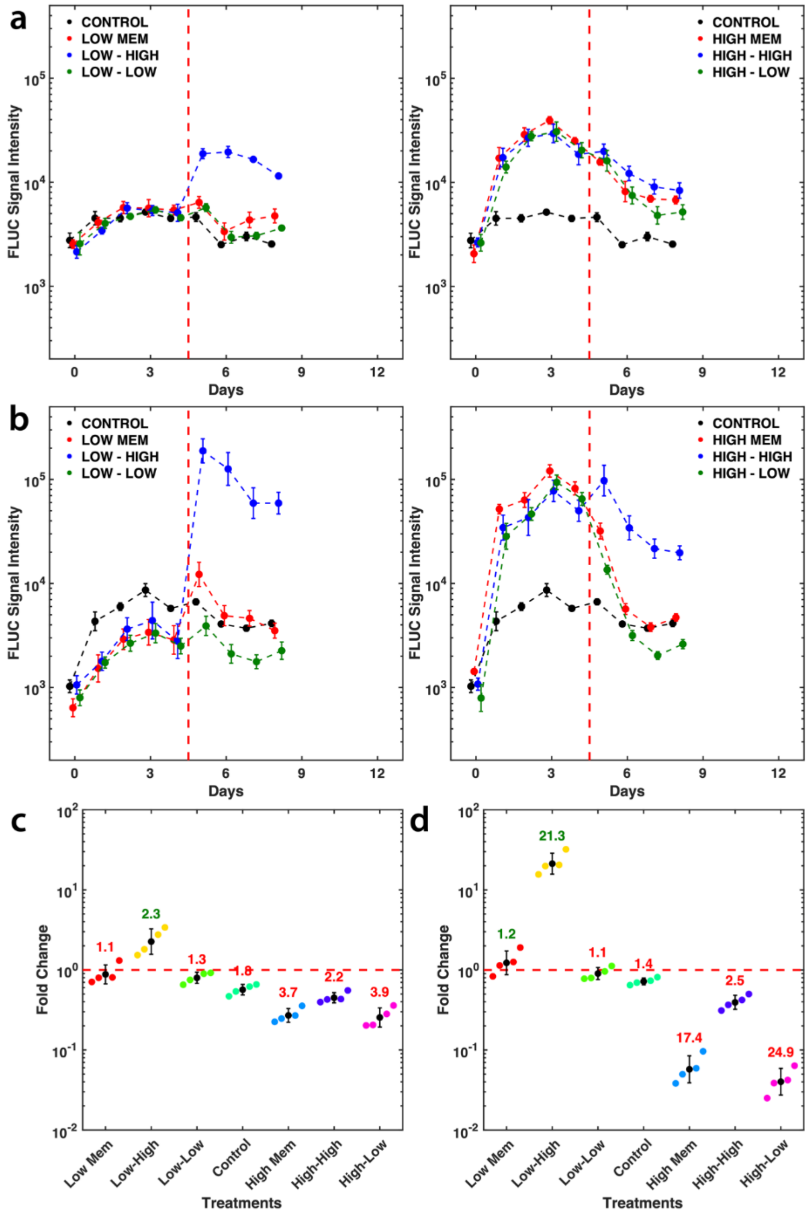


**Figure S4. Quantitative data analyses of Toggle 1.0 performance**. (**a-b**) Mean temporal luminescence levels of shoots (**a**) and roots (**b**) of plants under different treatments (See Materials and Methods for sample sizes). The x-axis shows the number of days on which luminescence data was collected starting from day 0. Data points on a same day are slightly shifted for clearer display. The y-axis shows the luminescence intensity. Error bars indicate standard errors. The red dashed line indicates the time when inducer was changed. (**c-d**) Average fold changes in luciferase activity of shoots (**c**) and roots (**d**) under different treatments (See Materials and Methods for sample sizes). Closed circles of different colors indicate fold change values of individual plants in corresponding treatment groups. The black closed circles represent the mean fold-change, and the black bar is the standard deviation. The number above each group of data-points is the average fold change of the corresponding group. Numbers in dark green represent increases in luminescence, while numbers in red indicate decreases. The red dashed line marks a fold change of one, *i.e.*, no change. All treatments are defined in detail in the Materials and Methods and Figure S3 legend.

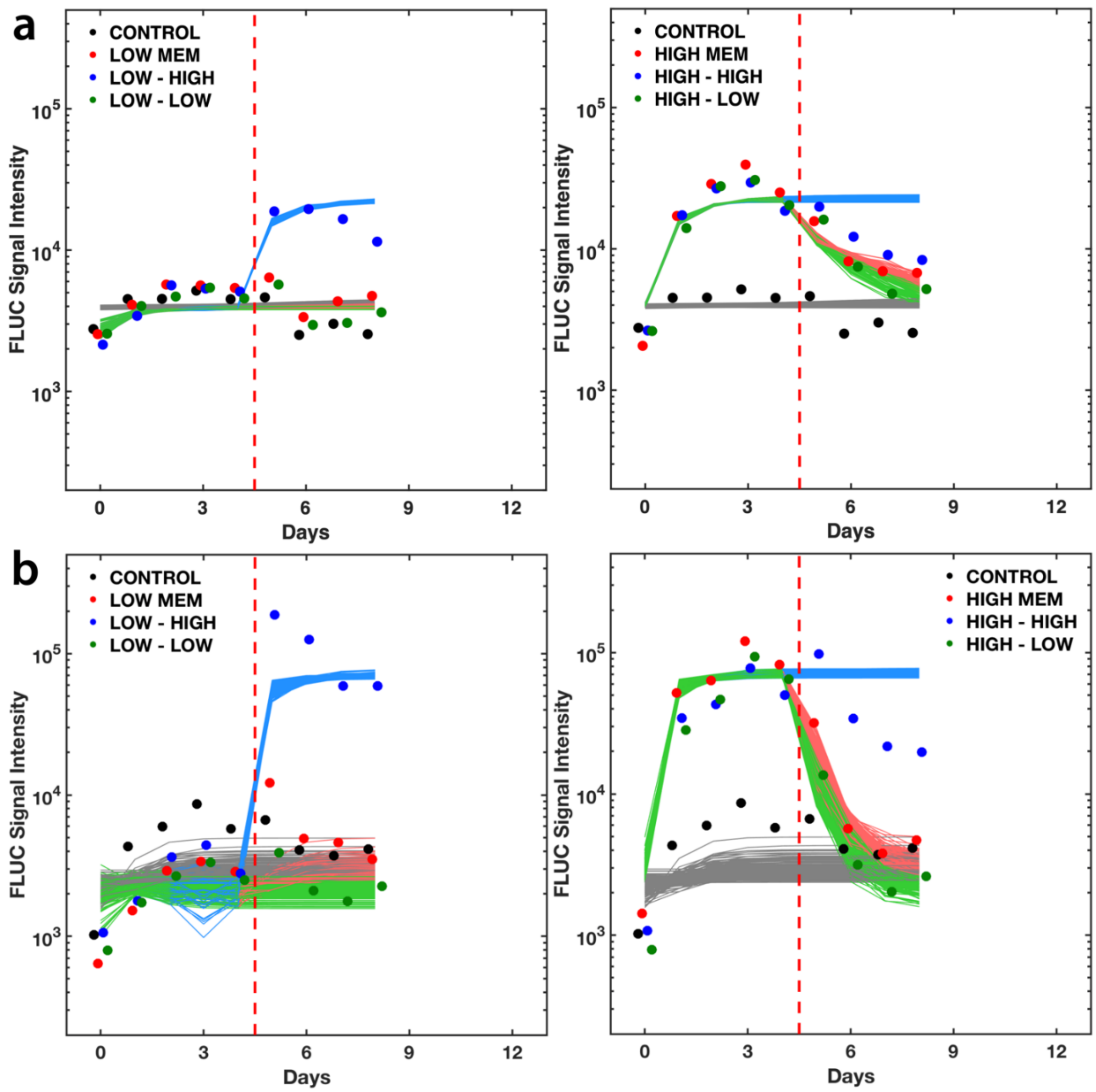


**Figure S5**. **Parameter calculation results of Toggle 1.0 using MCMC method**. Closed circles of different colors indicate mean luciferase intensity levels of shoots (**a**) and roots (**b**) of plants under different treatments from the experimental results shown in Figure S4a & b. Data points on a same day are shifted a little for clearer display. Each curve represents numerical solutions of ODEs using one estimated parameter set. Different colors correspond to different treatment conditions that are described in the legend to Figure S3. Fits plotted are best parameter sets as defined in *SI Materials and Methods*.

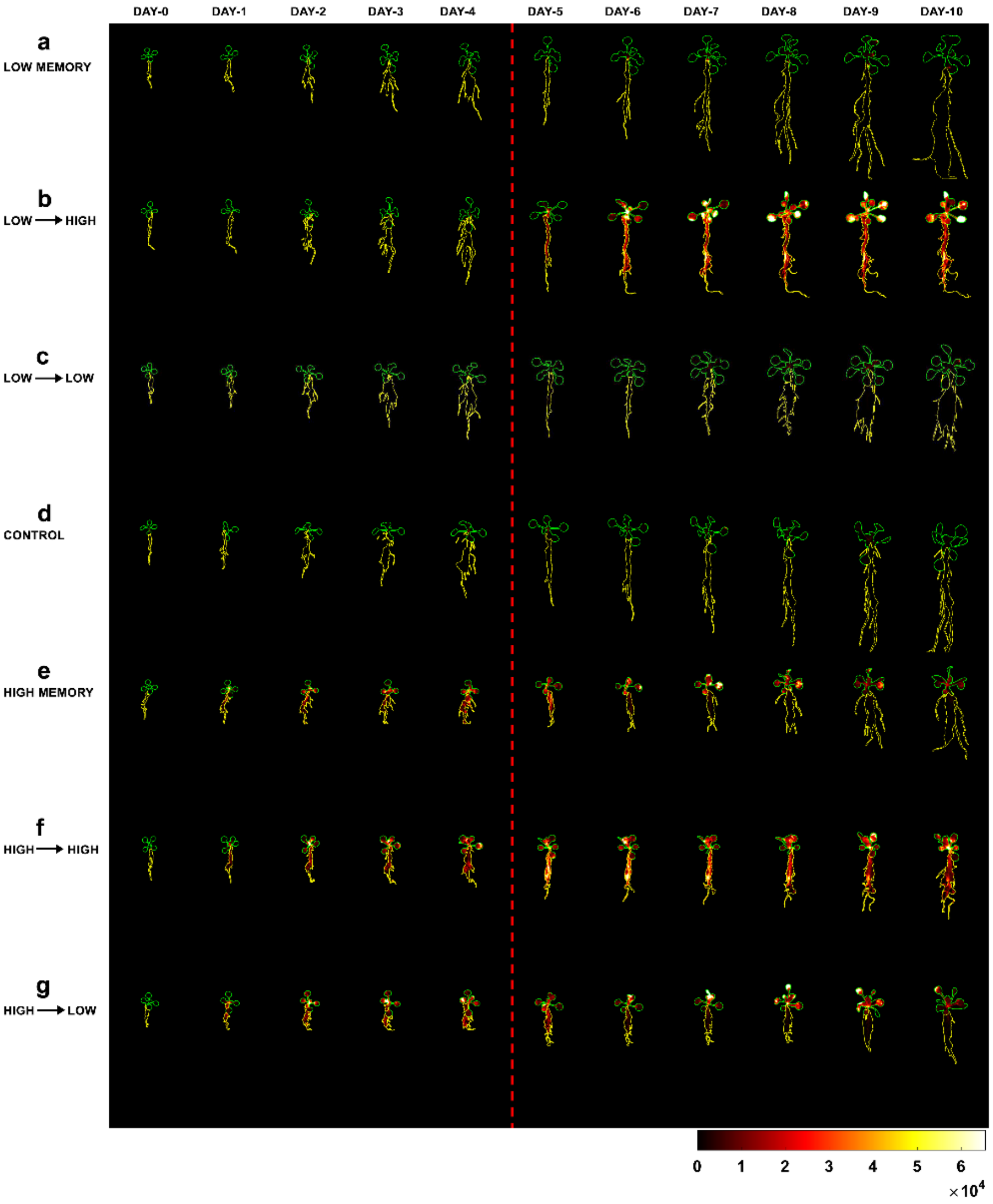


**Figure S6. Luminescence heat maps (linear scale) of selected plants containing Toggle 2.0 under different treatment conditions**. Green lines indicate ROIs drawn for the shoots, whereas yellow lines indicate those for the roots. All images use the same fire scale to indicate false-colored luminescence intensity, with black representing zero and white representing saturation according to the scale shown at the bottom. The colors are comparable and reflect luminescence intensity changes between different treatments. Inducer conditions were changed after imaging on Day 4, which is indicated by the red dashed line. (**a**) ***Low Memory***, plants incubated with DEX inducer for 4 days to establish the low state, then moved to media with no inducer to test for stability (memory) of the low state. (**b**) ***Low → High***, plants incubated with DEX inducer for 4 days to establish the low state, then moved to media with 4-OHT inducer to switch the circuit to the high state. (**c**) ***Low → Low***, plants incubated with DEX inducer throughout the experiment (low state). (**d**) ***Control***, plants incubated without inducer throughout the experiment. (**e**) ***High Memory***, plants incubated with 4-OHT inducer for 4 days to establish the high state, then moved to media with no inducer to test for stability (memory) of the high state. (**f**) ***High → High***, plants incubated with 4-OHT throughout the experiment (high state). (**g**) ***High → Low***, plants incubated with 4-OHT inducer for 4 days to establish the high state, then moved to media with DEX inducer to switch the circuit to the low state.

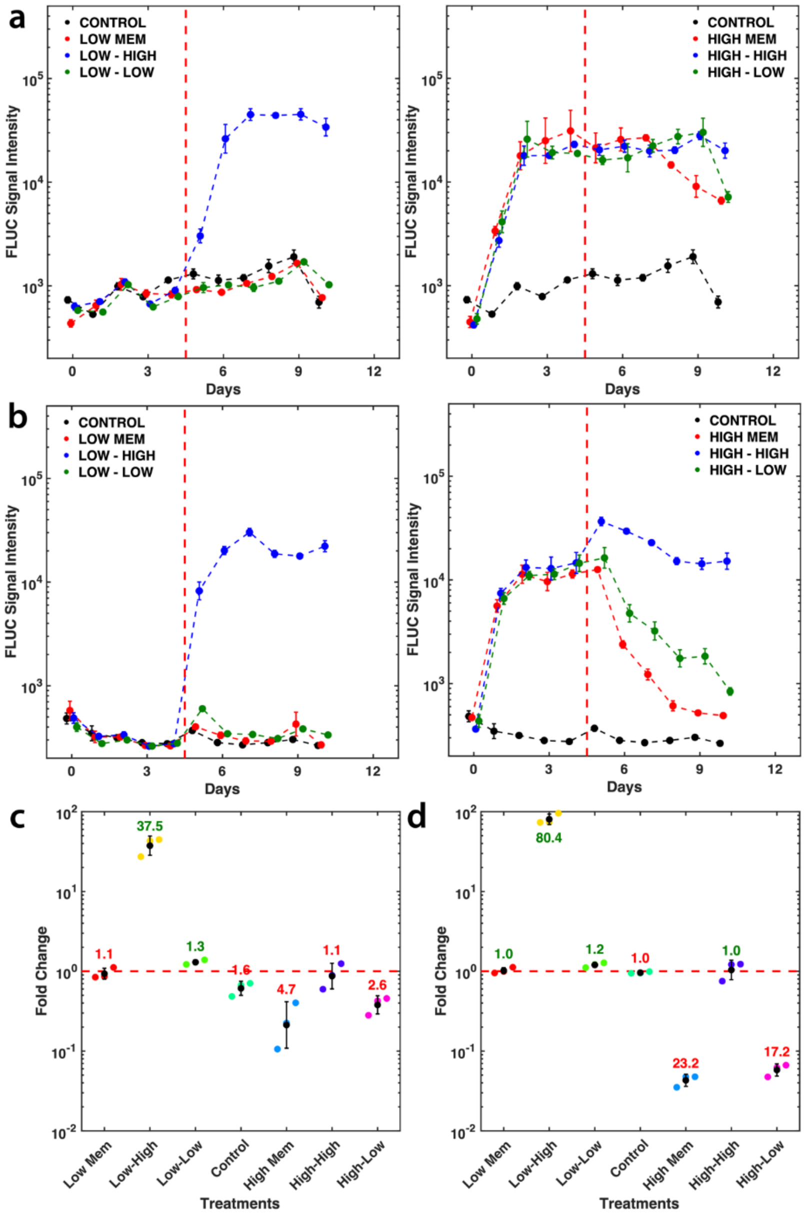


**Figure S7. Quantitative data analyses of Toggle 2.0 performance**. (**a-b**) Mean temporal luminescence levels of shoots (**a**) and roots (**b**) of plants under different treatments (See Materials and Methods for sample sizes). The x-axis shows the number of days on which luminescence data was collected starting from day 0. Data points on a same day are shifted a little for clearer display. The y-axis shows the luminescence intensity. Error bars indicate standard errors. The red dashed line indicates the time when inducer was changed. (**c-d**) Average fold changes of shoots (**c**) and roots (**d**) under different treatments (See Materials and Methods for sample sizes). Closed circles of different colors indicate fold-change values of individual plants in corresponding treatment groups. The black closed circles represent the mean fold change and the black bar the standard deviation. The number above each group of data-points is the average fold change of the corresponding group. Numbers in dark green represent increases in luminescence, while numbers in red indicate decreases. The red dashed line marks a fold change of one, *i.e.*, no change. All treatments are defined in detail in the Materials and Methods and Figure S6 legend.

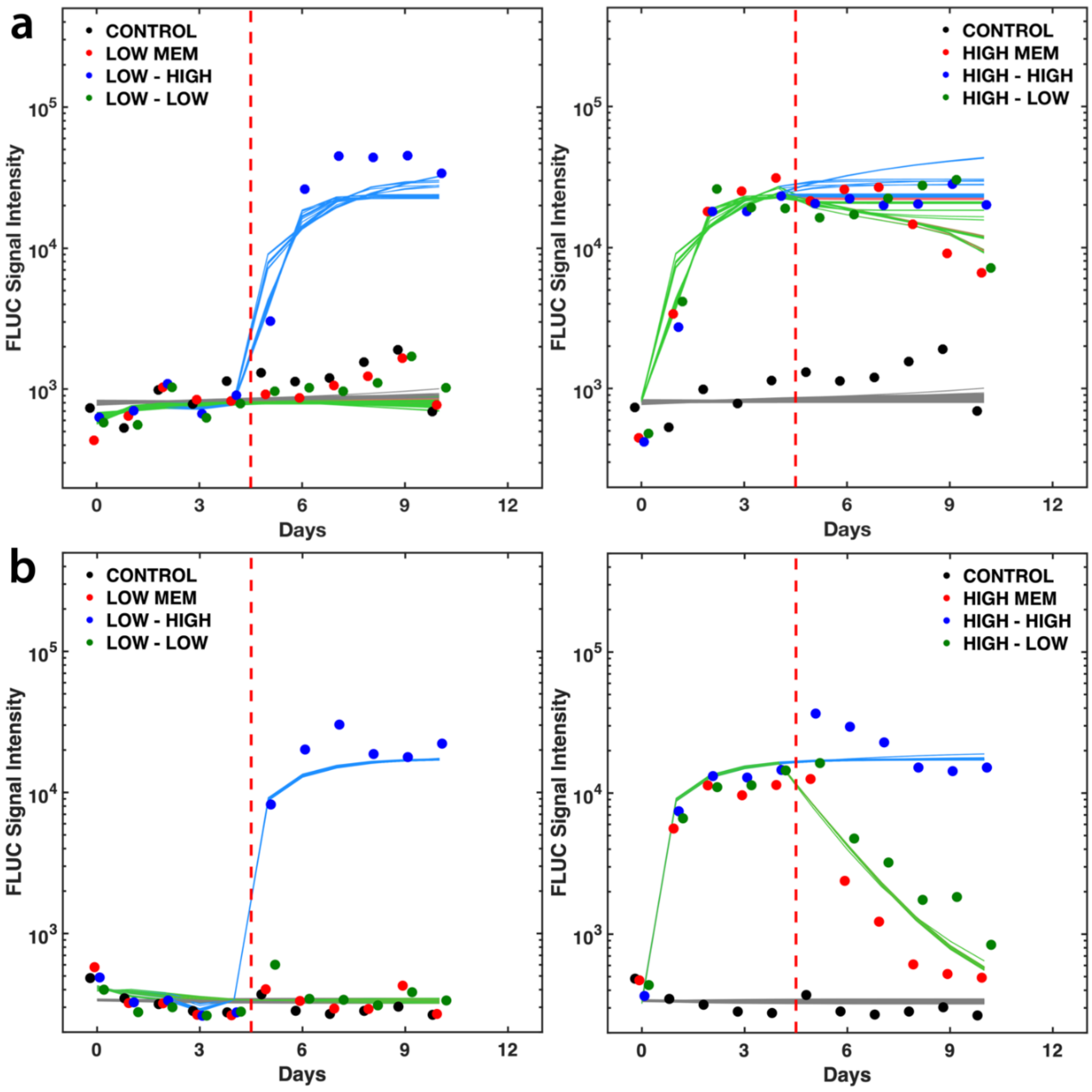


**Figure S8. Parameter calculation results of Toggle 2.0 using MCMC method**. Closed circles of different colors indicate mean luciferase intensity levels of shoots (**a**) and roots (**b**) of plants under different treatments from the experimental results shown in Figure S7a & b. Data points on a same day are slightly shifted for clearer display. Each curve represents numerical solutions of ODEs using one estimated parameter set. Different colors correspond to different treatment conditions that are described in the legend to Figure S6. Fits plotted are best parameter sets as defined in *SI Materials and Methods*.

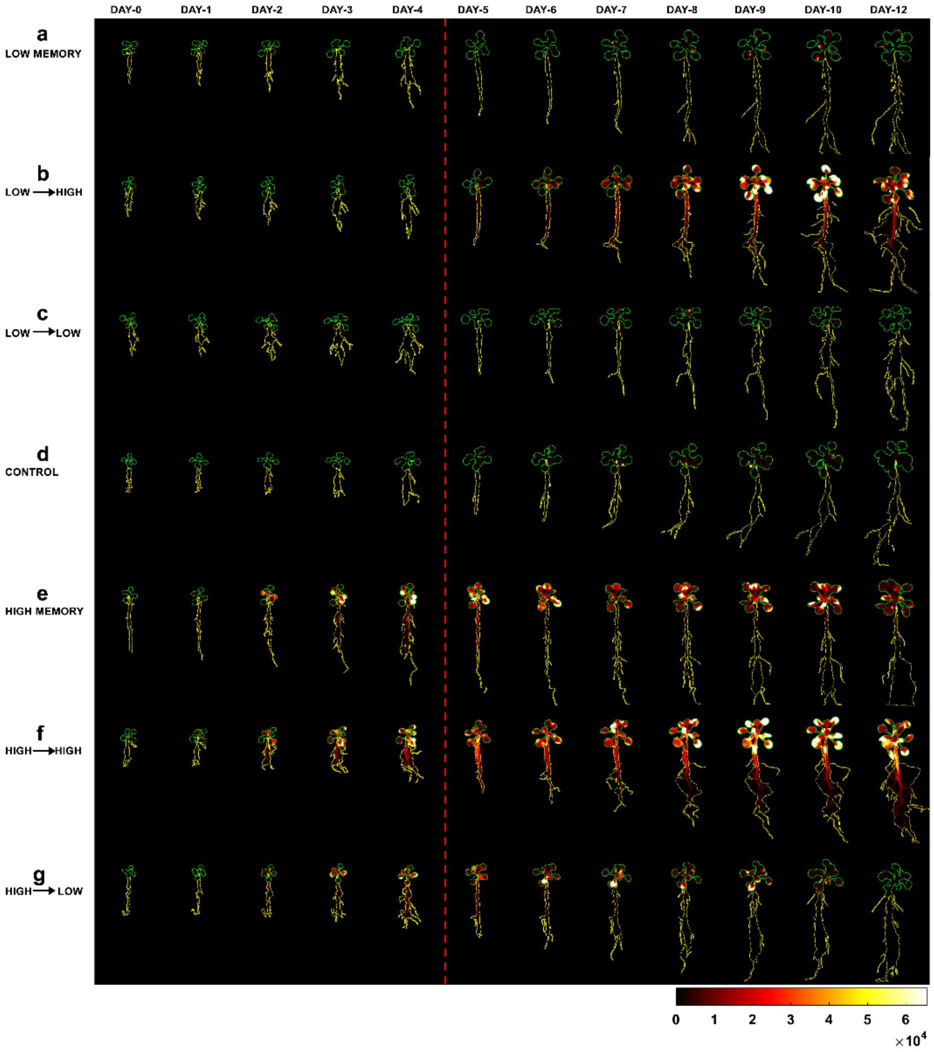


**Figure S9. Luminescence heat maps (linear scale) of selected plants showing Toggle 2.1 behavior under different treatments.** Green lines indicate ROIs drawn for the shoots, whereas yellow lines indicate those for the roots. All images use the same fire scale to indicate false-colored luminescence intensity, with black representing zero and white representing saturation according to the scale shown at the bottom. The colors reflect luminescence intensity changes between different treatments. Inducer conditions were changed after imaging on Day 4, which is indicated by the red dashed line. (**a**) ***Low Memory***, plants incubated with DEX inducer for 4 days to establish the low state, then moved to media with no inducer to test for stability (memory) of the low state. (**b**) ***Low → High***, plants incubated with DEX inducer for 4 days to establish the low state, then moved to media with 4-OHT inducer to switch the circuit to the high state. (**c**) ***Low → Low***, plants incubated with DEX inducer throughout the experiment (low state). (**d**) ***Control***, plants incubated without inducer throughout the experiment. (**e**) ***High Memory***, plants incubated with 4-OHT inducer for 4 days to establish the high state, then moved to media with no inducer to test for stability (memory) of the high state. (**f**) ***High → High***, plants incubated with 4-OHT throughout the experiment (high state). (**g**) ***High → Low***, plants incubated with 4-OHT inducer for 4 days to establish the high state, then moved to media with DEX inducer to switch the circuit to the low state. See Figure 6, main text, for brightfield photos of plants.

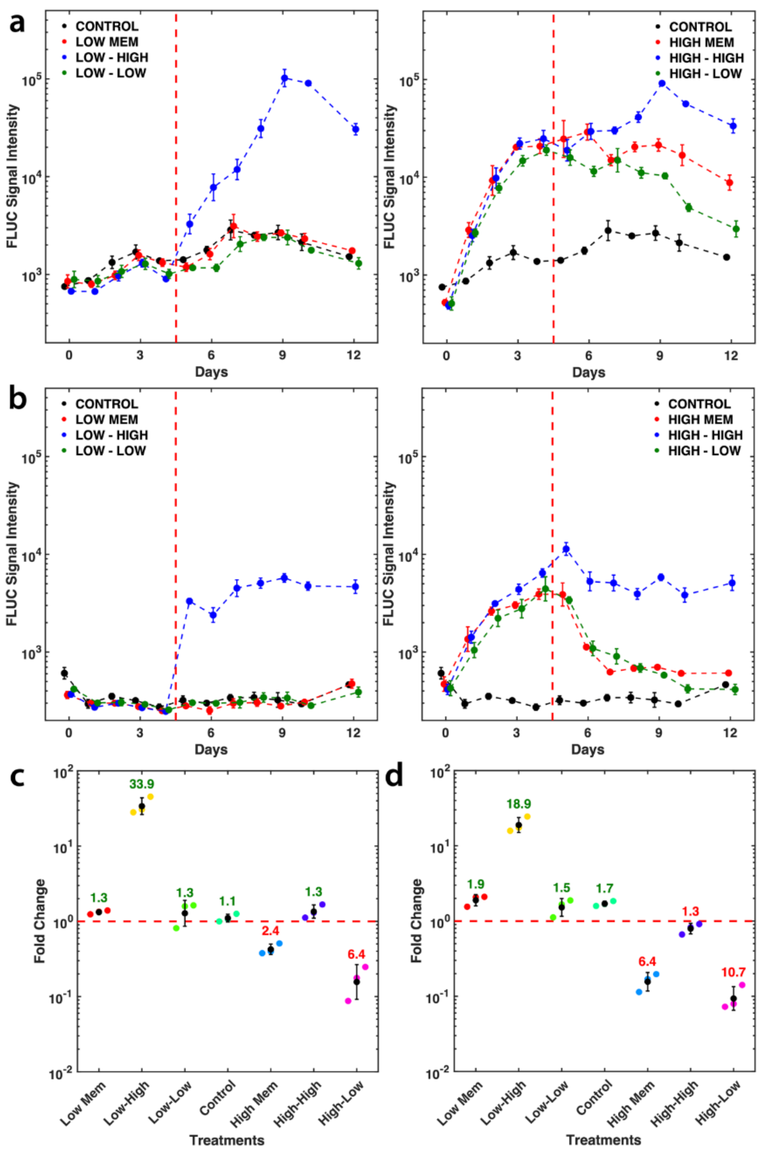


**Figure S10. Quantitative data analyses of Toggle 2.1 performance**. (**a-b**) Mean temporal luminescence levels of shoots (**a**) and roots (**b**) of plants under different treatments (See Materials and Methods for sample sizes). The x-axis shows the number of days in which luminescence data was collected starting from day 0. Data points on a same day are slightly shifted for clearer display. The y-axis shows the luminescence intensity. Error bars indicate standard errors. The red dashed line indicates the time when inducer was changed, or media was refreshed. (**c-d**) Average fold changes of shoots (**c**) and roots (**d**) under different treatments (See Materials and Methods for sample sizes). Closed circles of different colors indicate fold-change values of individual plants under corresponding treatments. The black closed circles indicate the mean fold-change and the black bars the standard deviations. Numbers above the groups are the average fold-change of the corresponding treatment. Numbers in dark green colored number indicate increases in luciferase level and red colored numbers decreases. Red dashed line marks the baseline of one, indicating no change. All treatments were defined in detail in the Materials and Methods and Figure S9 legend.

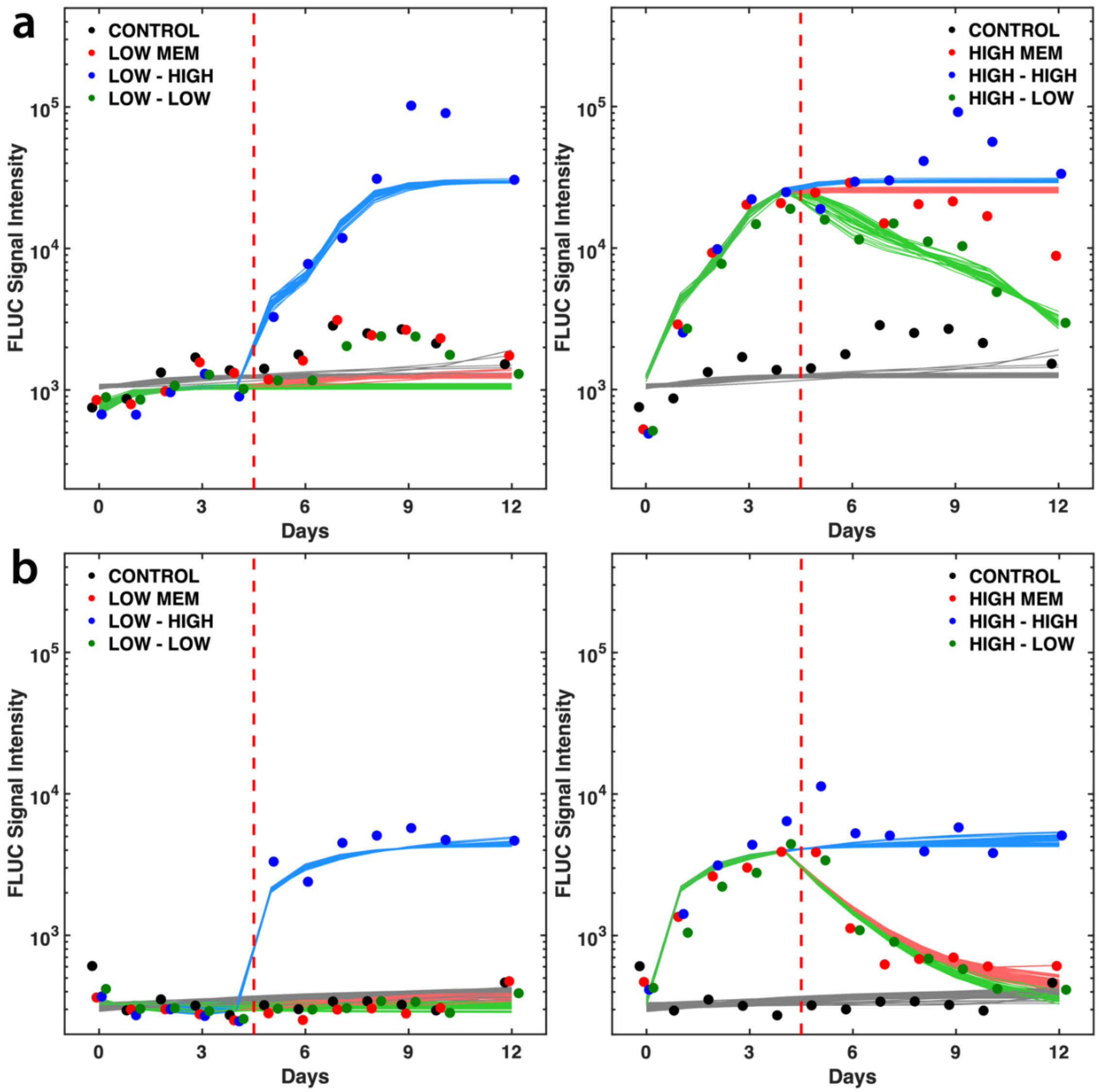


**Figure S11. Parameter calculation results of Toggle 2.1 using MCMC method**. Parameter values estimated by MCMC method fitted to experimental data of shoots (**a**) and roots (**b**) of plants under different treatments from the experimental results shown in Figure S10a & b. Data points on a same day are slightly shifted for clearer display. Each curve represents numerical solutions of ODEs using one estimated parameter set. The High to Low (green curve in right panel in (**a**)) shows the improvement in switching dynamics of Toggle 2.1 compared to Toggle 2.0 as shown in Figure S8a. Different colors correspond to different treatment conditions that are described in the legend to Figure S9. Fits plotted are best parameter sets as defined in *SI Materials and Methods*.

**a**

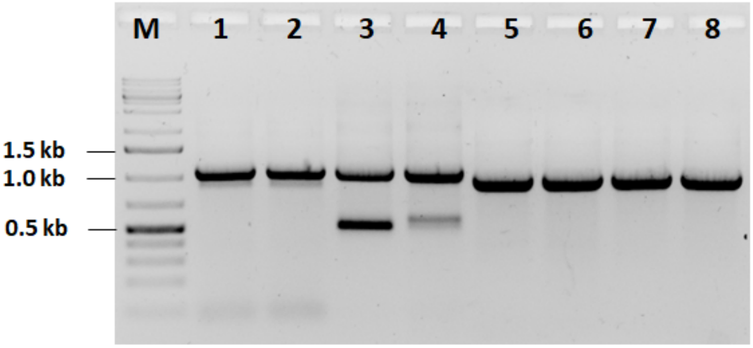


**b**

AGAAGCAGAGACTGTGAAGATGAGATTGCGTTGTCTAGCTTCAGGAATGGTGGAGAAGG
GTATCGAAGTGTAGATATGTGGTCTGGTTAAGGGTAATAATGGAAGAGAAAGTGAAGAA
GATGAAGCAATTTCTATGTACATCTTACACAAAGGATTTTTTAAAAACAAAAGTACATTA
AATTTTAGTTTCCAGCCTCATCGACAGTCTATTTATAAGTAATATATCAGAGTTGAGAGTC
AGCAAAATTTAAGCTCATCTCTATCTCTCAAAAGCAAAGTAAAGCTTAACCAATGGAGAT
AAGGTATTTTCATATCCGTAGTTACGTACAATTCGTAAAATGTGTTTTCGTACCTAAATAT
ATTTGTAGAGTAAACTATAACCCAAACCAAAAAAATATTTAAATTTCATATCTAAAATTTG
TGCAAAAACAAGAGTCAAATTAATTAGTGTTAGCCAACATTGTTACAGTGCTAAGTTAAG
CGAACACATATATGATGATATGTCTTTAAAAGATATGTAGACTCGAACCTGTAAGTTATA
GCTTTTAGATAAGTATCTAATACCGATTTATCCGACAACCTTCTCCGGGTTATACACAATA
AAACAAATTTATAAATCTTAAATATCTGGTATCAATTATCAGTTCTTATTTACGGACCAAA
TTTCACGCTAAAATGAAAACCCACTATTGATAAATTCGTGGAAAAGATCAAATCAGAAGC
TAAGAACAATGACAAGACTCATTTAGATTCACTGCTTTATGTATTTTGTTAGCTATGTCTT
TTGAAATCGAAATGGAAGAAGAGTCTATTGAGAAGATGTATCAAATGATGGCGGAAATTT
TAATTGTTGTTGATCCTACATCAAAATCCTAGAATTTGATTATTTTTTCCAATTCACAACG
ACATTCTTTTGTTAATCTGAAAACACAACACACTTAAGGTATATAAACTAAAACGTTTGGG
TAAATCTCGATGAAGATTATGAGATAAAAATGGTTATCAGCTTGCATGCCGGTCGATCTA
GTAACATAGATGACACCGCGCGCGATAATTTATCCTAGTTTGCGCGCTATATTTTGTTTTC
TATCGCGTATTAAATGTATAATTGCGGGACTCTAATCATAAAAACCCATCTCATAAATAAC
GTCATGCATTACATGTTAATTATTACATGCTTAACGTAATTCAACAGAAATTATATGATAA
TCATCGCAAGACCGGCAACAGGATTCAATCTTAAGAAACTTTATTGCCAAATGTTTGAAC
GATCTGCTTGACTCTAGGGGTCATCAGATTTCGGTGACGGGCAGGACCGGACGGGGCGG
CACCGGCAGGCTGAAGTCCAGCTGCCAGAAACCCACGTCATG

**Figure S12. Determination of chromosomal location and copy number estimation of the transgene**. (**a**) Secondary and tertiary TAIL-PCR products of Toggle 2.0 resolved in a 1% agarose gel stained with ethidium bromide. PCR was performed using genomic DNA isolated from both shoot (Lanes 1, 2, 5 & 6) and root (Lanes 3, 4, 7 & 8) tissues. Samples 1-4 are secondary and samples 5-8 are tertiary TAIL-PCR products. The expected slight decrease in size of tertiary TAIL-PCR products is due to the use of nested primers. M, marker (GeneRuler 1-kb plus ladder, Thermo Scientific). (**b**) DNA sequence of Toggle 2.0 tertiary TAIL-PCR product cloned in the pJET vector. T-DNA insertion was found in Chromosome 5 in the intergenic between AT5G39980 and AT5G39990. Gray shaded region is from Arabidopsis chromosome 5, the unshaded sequence is part of the T-DNA border and T-NOS, and nucleotides in blue are the LB1 primer.

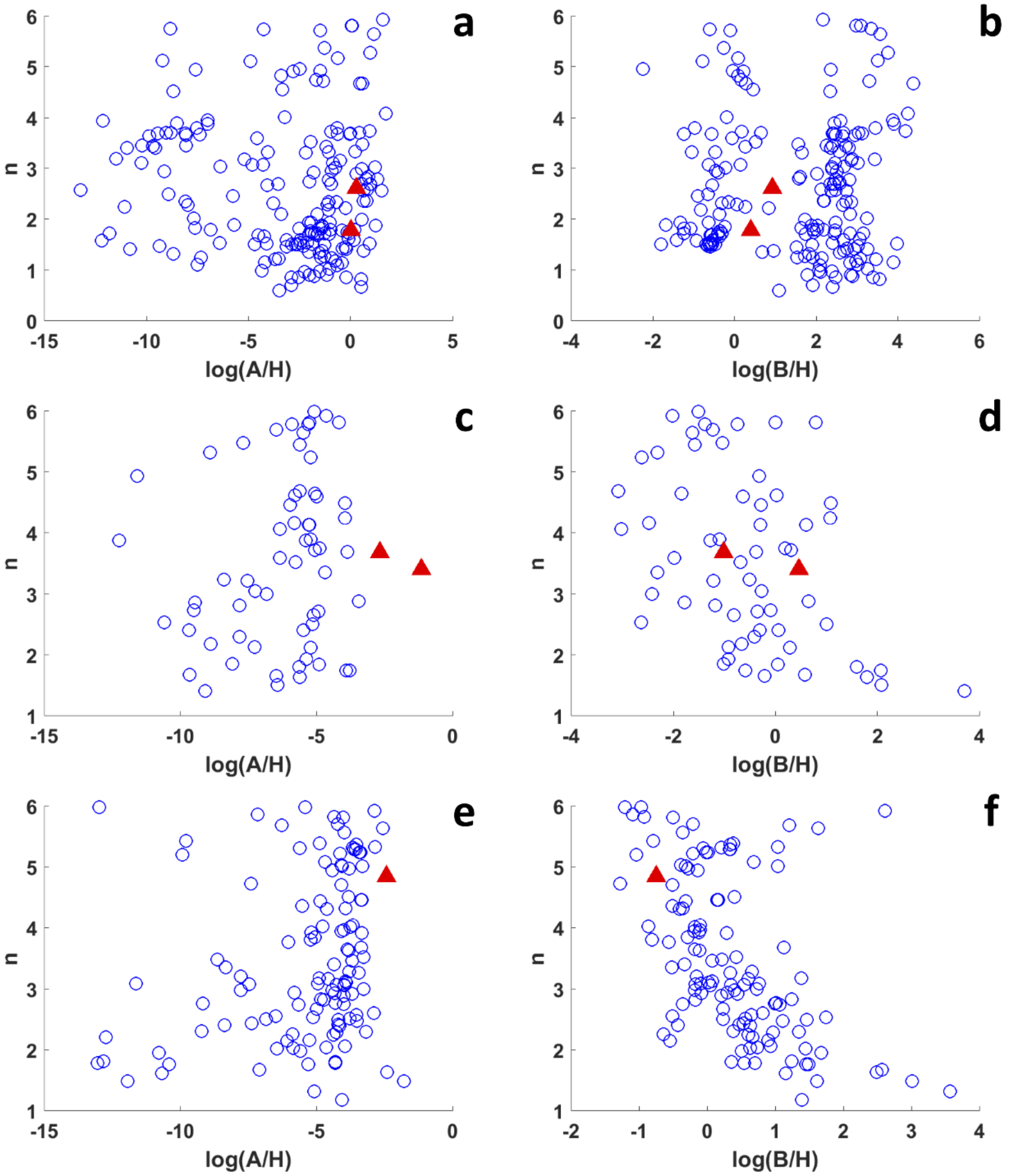


**Figure S13. Comparisons between dimensionless parameter values estimated from protoplast assays and whole plant luciferase data.** The x-axis is the natural logarithm of the dimensionless variables A/H or B/H, as defined in Equation S5. The rationale behind using these parameters for comparison of the results of the protoplast with whole plant data is explained in the SI Notes. The y-axis represents the values of Hill coefficients. The empty blue circles are parameter value combinations estimated from the luciferase levels of shoots quantified in this study and the filled red triangles the parameter values from protoplast assays published previously. (**a-b**) $NOS_{2xGal4}$.EAR; (**c-d**) $35S_{2xLexA}$.EAR; (**e-f**) $35S_{4xLexA}$TATA.OFPx.

**Table S1.** Sequence information of all the parts used to construct the toggle circuits.

| Name | Type | DNA sequence |
|---|---|---|
| 35S | Promoter | CATGGAGTCAAAGATTCAAATAGAGGACCTAACAGAACTCGCCGTAAAGACTGGC<br>GAACAGTTCATACAGAGTCTCTTACGACTCAATGACAAGAAGAAAATCTTCGTCA<br>ACATGGTGGAGCACGACACACTTGTCTACTCCAAAAATATCAAAGATACAGTCTC<br>AGAAGACCAAAGGGCAATTGAGACTTTTCAACAAAGGGTAATATCCGGAAACCTC<br>CTCGGATTCCATTGCCCAGCTATCTGTCACTTTATTGTGAAGATAGTGGAAAAGGA<br>AGGTGGCTCCTACAAATGCCATCATTGCGATAAAGGAAAGGCCATCGTTGAAGAT<br>GCCTCTGCCGACAGTGGTCCCAAAGATGGACCCCCACCCACGAGGAGCATCGTGG<br>AAAAAGAAGACGTTCCAACCACGTCTTCAAAGCAAGTGGATTGATGTGATATCTC<br>CACTGACGTAAGGGATGACGCACAATCCCACTATCCTTCGCAAGACCCTTCCTCTA<br>TATAAGGAAGTTCATTTCATTTGGAGAGAACACGGGGGACTCTCC |
| FMV | Promoter | GGCCGCAGGATTTAGCAGCATTCCAGATTGGGTTCAATCAACAAGGTACGAGCCA<br>TATCACTTTATTCAAATTGGTATCGCCAAAACCAAGAAGGAACTCCCATCCTCAAA<br>GGTTTGTAAGGAAGAATTCTCAGTCCAAAGCCTCAACAAGGTCAGGGTACAGAGT<br>CTCCAAACCATTAGCCAAAAGCTACAGGAGATCAATGAAGAATCTTCAATCAAAG<br>TAAACTACTGTTCCAGCACATGCATCATGGTCAGTAAGTTTCAGAAAAAGACATC<br>CACCGAAGACTTAAAGTTAGTGGGCATCTTTGAAAGTAATCTTGTCAACATCGAG<br>CAGCTGGCTTGTGGGGACCAGACAAAAAAGGAATGGTGCAGAATTGTTAGGCGC<br>ACCTACCAAAAGCATCTTTGCCTTTATTGCAAAGATAAAGCAGATTCCTCTAGTAC<br>AAGTGGGGAACAAAATAACGTGGAAAAGAGCTGTCCTGACAGCCCACTCACTAAT<br>GCGTATGACGAACGCAGTGACGACCACAAAAGAATTCCCTCTATATAAGAAGGCA<br>TTCATTCCCATTTGAAGGATCATCAGATACTCAACC |
| $35S_{2xLexA}$ | Synthetic promoter | TGAGACTTTTCAACAAAGGGTAATATCGGGAAACCTCCTCGGATTCCATTGCCCA<br>GCTATCTGTCACTTCATCAAAAGGACAGTAGAAAAGGAAGGTGGCACCTACAAAT<br>GCCATCATTGCGATAAAGGAAAGGCTATCGTTCAAGATGCCTCTGCCGACAGTGG<br>TCCCAAAGATGGACCCCCACCCACGAGGAGCATCGTGGAAAAAGAAGACGTTCC<br>AACCACGTCTTCAAAGCAAGTGGATTGATGTGATATCTCCACTGACGTAAGGGAT<br>GACGCACAATCCCACTATCCTTCGCAAGACCCTTCCTCTATATAAGGAAGTTCATT<br>TCATTTGGAGAGGAACGCGTACTGTACATATAACCACTGGTTTTATATACAGCAGT |
| $35S_{4xLexA}$TATA | Synthetic promoter | TGAGACTTTTCAACAAAGGGTAATATCGGGAAACCTCCTCGGATTCCATTGCCCA<br>GCTATCTGTCACTTCATCAAAAGGACAGTAGAAAAGGAAGGTGGCACCTACAAAT<br>GCCATCATTGCGATAAAGGAAAGGCTATCGTTCAAGATGCCTCTGCCGACAGTGG<br>TCCCAAAGATGGACCCCCACCCACGAGGAGCATCGTGGAAAAAGAAGACGTTCC<br>AACCACGTCTTCAAAGCAAGTGGATTGATGTGATATCTCCACTGACGTAAGGGAT<br>GACGCACAATCCCACTATCCTTCGCAAGACCCTTCCTCACTGTACATATAACCACT<br>GGTTTTATATACAGCAGTACTGTACATATAACCACTGGTTTTATATACAGCAGTTA<br>TATAAGGAAGTTCATTTCATTTGGAGAGGA |
| $NOS_{2xGal4}$ | Synthetic promoter | AGGCGGGAAACGACAATCTGATCATGAGCGGAGAATTAAGGGAGTCACGTTATG<br>ACCCCCGCCGATGACGCGGGACAAGCCGTTTTACGTTTGGAACTGACAGAACCGC<br>AACGATTGAAGGAGCCACTCAGCCGCGGGTTTCTGGAGTTTAATGAGCTAAGCAC<br>ATACGTCAGAAACCATTATTGCGCGTTCAAAAGTCGCCTAAGGTCACTATCAGCT<br>AGCAAATATTTCTTGTCAAAAATGCTCCACTGACGTTCCATAAATTCCCCTCGGTA<br>TCCAATTAGAGTCTCATATTCACTCTCAATCCAAATAATCTGCACCGACGCGTCGG<br>AAGACTCTCCTCCGAGCGGAAGACTCTCCTCCG |

| | | |
|---|---|---|
| GEAR | Synthetic Repressor | ATGAAGTTGTTAAGTTCTATTGAACAGGCTTGTGATATTTGTAGATTGAAGAAGCTCAAGTGTAGTAAAGAGAAACCTAAGTGCGCTAAGTGTCTTAAAAATAACTGGGAGTGCAGATATTCTCCTAAGACTAAGAGATCACCTCTTACTAGGGCTCATCTCACAGAGGTGGAGTCTAGGCTTGAGAGATTGGAGCAGTTGTTCCTTTTGATTTTTCCAAGAGAAGATCTTGATATGATTCTTAAGATGGATTCTCTTCAAGATATTAAGGCTCTTCTTACTGGTTTATTCGTTCAAGATAATGTGAATAAGGATGCAGTGACTGATAGACTTGCATCAGTTGAAACTGATATGCCTCTTACATTAAGACAACACAGGATTTCTGCTACTTCAAGTTCTGAAGAAAGTTCTAATAAGGGTCAAAGGCAATTAACCGTTTCTCTTGATTTGGACCTCGAGCTTCGTTTGGGATTCGCTTGA |
| LEAR | Synthetic Repressor | ATGAAAGCTCTTACTGCTAGACAACAAGAAGTTTTTGATTTGATTAGGGATCATATTTCTCAGACAGGTATGCCTCCTACTAGAGCTGAGATCGCTCAGAGACTCGGTTTCAGATCTCCTAACGCTGCTGAAGAGCATCTTAAGGCTCTTGCTAGAAAGGGAGTTATTGAGATTGTGAGTGGAGCATCAAGAGGTATTAGGTTGCTTCAAGAGGAAGAAGAGGGACTTCCTCTTGTTGGTAGAGTTGCAGCTGGTGAGCCTCTTGATTTGGACCTCGAGCTTCGTTTGGGATTCGCTTGA |
| LOFPx | Synthetic Repressor | ATGAAAGCTCTTACTGCTAGACAACAAGAAGTTTTTGATTTGATTAGGGATCATATTTCTCAGACAGGTATGCCTCCTACTAGAGCTGAGATCGCTCAGAGACTCGGTTTCAGATCTCCTAACGCTGCTGAAGAGCATCTTAAGGCTCTTGCTAGAAAGGGAGTTATTGAGATTGTGAGTGGAGCATCAAGAGGTATTAGGTTGCTTCAAGAGGAAGAAGAGGGACTTCCTCTTGTTGGTAGAGTTGCAGCTGGTGAGCCTAGCTTGGCGGTGGTGAAGAAGTCGGTGGATCCAAACAAAGATTTCAGGGAATCAATGGTGGAGATGATAGCAGAGAACAAGATAAGAGCATCAAATGACCTAGAAGAGCTTCTTGCTTGCTACCTTTCGTTAAATCCAAAGGAATATCACGATCTTATTATCAAAGTGTTCGAACAAATCTGGCTTGAACTGATAAACCCA |
| NOST | Terminator | CGTTCAAACATTTGGCAATAAAGTTTCTTAAGATTGAATCCTGTTGCCGGTCTTGCGATGATTATCATATAATTTCTGTTGAATTACGTTAAGCATGTAATAATTAACATGTAATGCATGACGTTATTTATGAGATGGGTTTTTATGATTAGAGTCCCGCAATTATACATTTAATACGCGATAGAAAACAAAATATAGCGCGCAAACTAGGATAAATTATCGCGCGCGGTGTCATCTATGTTACTAGATCGGGCGGG |
| OCST | Terminator | GGAGCTGCTTTAATGAGATATGCGAGACGCCTATGATCGCATGATATTTGCTTTCAATTCTGTTGTGCACGTTGTAAAAAACCTGAGCATGTGTAGCTCAGATCCTTACCGCCGGTTTCGGTTCATTCTAATGAATATATCACCCGTTACTATCGTATTTTTATGAATAATATTCTCCGTTCAATTTACTGATTGTACCCTACTACTTATATGTACAATATTAAAATGAAAACAATATATTGTGCTGAATAGGTTTATAGCGACATCTATGATAGAGCGCCACAATAACAAACAATTGCGTTTTATTATTACAAATCCAATTTTAAAAAAAGCGGCAGAACCGGTCAAACCTAAAAGACTGATTACATAAATCTTATTCAAATTTCAAAAGGCCCCAGGGGCTAGTATCTACGACACACCGAGCGGCGAACTAATAACGTTCACTGAAGGGAACTCCGGTTCCCCGCCGGCGCGCATGGGTGAGATTCCTTGAAGTTGAGTATTGGCCGTCCGCTCTACCGAAAGTTACGGGCACCATTCAACCCGGTCCAGCACGGCGGCCGGGTAACCGACTTGCTGCCCCGAGAATTATGCAGCATTTTTTTGGTGTATGT |

| | | |
|---|---|---|
| Pea3AT | Terminator | GCTTCGGGCCTCCCAGCTTTCGTCCGTATCATCGGTTTCGACAACGTTCGTCAAGT<br>TCAATGCATCAGTTTCATTGCCCACACACCAGAATCCTACTAAGTTTGAGTATTAT<br>GGCATTGGAAAAGCTGTTTTCTTCTATCATTTGTTCTGCTTGTAATTTACTGTGTTC<br>TTTCAGTTTTTGTTTTCGGACATCAAAATGCAAATGGATGGATAAGAGTTAATAAA<br>TGATATGGTCCTTTTGTTCATTCTCAAATTATTATTATCTGTTGTTTTTACTTTAATG<br>GGTTGAATTTAAGTAAGAAAGGAACTAACAGTGTGATATTAAGGTGCAATGTTAG<br>ACATATAAAACAGTCTTTCACCTCTCTTTGGTTATGTCTTGAATTGGTTTGTTTCTT<br>CACTTATCTGTGTAATCAAGTTTACTATGAGTCTATGATCAAGTAATTATGCAATC<br>AAGTTAAGTACAGTATAGGCTTT |
| E9T | Terminator | ATTATGGCATTGGGAAAACTGTTTTTCTTGTACCATTTGTTGTGCTTGTAATTTACT<br>GTGTTTTTTATTCGGTTTTCGCTATCGAACTGTGAAATGGAAATGGATGGAGAAGA<br>GTTAATGAATGATATGGTCCTTTTGTTCATTCTCAAATTAATATTATTTGTTTTTTC<br>TCTTATTTGTTGTGTGTTGAATTTGAAATTATAAGAGATATGCAAACATTTTGTTTT<br>GAGTAAAAATGTGTCAAATCGTGGCCTCTAATGACCGAAGTTAATATGAGGAGTA<br>AAACATCCCAAAC |
| TB | Transcription block | AATAAAATATCTTTATTTTCATTACATCTGTGTGTTGGTTTTTTGTGTGAATCGATA<br>GTACTAACATACGCTCTCCATCAAAACAAAACGAAACAAAACAAACTAGCAAAAT<br>AGGCTGTCCCCAGTGCAAGTGCAGGTGCCAGAACATTTCTCT |
| F-LUC | Gene | ATGGAAGACGCCAAAAACATAAAGAAAGGCCCGGCGCCATTCTATCCTCTAGAGG<br>ATGGAACCGCTGGAGAGCAACTGCATAAGGCTATGAAGAGATACGCCCTGGTTCC<br>TGGAACAATTGCTTTTACAGATGCACATATCGAGGTGAACATCACGTACGCGGAA<br>TACTTCGAAATGTCCGTTCGGTTGGCAGAAGCTATGAAACGATATGGGCTGAATA<br>CAAATCACAGAATCGTCGTATGCAGTGAAAACTCTCTTCAATTCTTTATGCCGGTG<br>TTGGGCGCGTTATTTATCGGAGTTGCAGTTGCGCCCGCGAACGACATTTATAATGA<br>ACGTGAATTGCTCAACAGTATGAACATTTCGCAGCCTACCGTAGTGTTTGTTTCCA<br>AAAAGGGGTTGCAAAAAATTTTGAACGTGCAAAAAAAATTACCAATAATCCAGA<br>AAATTATTATCATGGATTCTAAAACGGATTACCAGGGATTTCAGTCGATGTACACG<br>TTCGTCACATCTCATCTACCTCCCGGTTTTAATGAATACGATTTTGTACCAGAGTCC<br>TTTGATCGTGACAAAACAATTGCACTGATAATGAATTCCTCTGGATCTACTGGGTT<br>ACCTAAGGGTGTGGCCCTTCCGCATAGAACTGCCTGCGTCAGATTCTCGCATGCCA<br>GAGATCCTATTTTTGGCAATCAAATCATTCCGGATACTGCGATTTTAAGTGTTGTT<br>CCATTCCATCACGGTTTTGGAATGTTTACTACACTCGGATATTTGATATGTGGATTT<br>CGAGTCGTCTTAATGTATAGATTTGAAGAAGAGCTGTTTTTACGATCCCTTCAGGA<br>TTACAAAATTCAAAGTGCGTTGCTAGTACCAACCCTATTTTCATTCTTCGCCAAAA<br>GCACTCTGATTGACAAATACGATTTATCTAATTTACACGAAATTGCTTCTGGGGGC<br>GCACCTCTTTCGAAAGAAGTCGGGGAAGCGGTTGCAAAACGCTTCCATCTTCCAG<br>GGATACGACAAGGATATGGGCTCACTGAGACTACATCAGCTATTCTGATTACACC<br>CGAGGGGGATGATAAACCGGGCGCGGTCGGTAAAGTTGTTCCATTTTTTGAAGCG<br>AAGGTTGTGGATCTGGATACCGGGAAAACGCTGGGCGTTAATCAGAGAGGCGAAT<br>TATGTGTCAGAGGACCTATGATTATGTCCGGTTATGTAAACAATCCGGAAGCGAC<br>CAACGCCTTGATTGACAAGGATGGATGGCTACATTCTGGAGACATAGCTTACTGG<br>GACGAAGACGAACACTTCTTCATAGTTGACCGCTTGAAGTCTTTAATTAAATACAA<br>AGGATATCAGGTGGCCCCCGCTGAATTGGAATCGATATTGTTACAACACCCCAAC<br>ATCTTCGACGCGGGCGTGGCAGGTCTTCCCGACGATGACGCCGGTGAACTTCCCG<br>CCGCCGTTGTTGTTTTGGAGCACGGAAAGACGATGACGGAAAAAGAGATCGTGGA<br>TTACGTCGCCAGTCAAGTAACAACCGCGAAAAAGTTGCGCGGAGGAGTTGTGTTT<br>GTGGACGAAGTACCGAAAGGTCTTACCGGAAAACTCGACGCAAGAAAAATCAGA<br>GAGATCCTCATAAAGGCCAAGAAGGGCGGAAAGTCCAAATTGTAA |

| | | |
|---|---|---|
| 10xN1 | 4-OHT Inducible Promoter | GGGGTAGAAAAAGGGGTAGAAAACCAAGGGGTAGAAAAAGGGGTAGAAAACCA AGGGGTAGAAAAAGGGGTAGAAAACCAAGGGGTAGAAAAAGGGGTAGAAAACC AAGGGGTAGAAAAAGGGGTAGAAGCAAGACCCTTCCTCTATATAAGGAAGTTCAT TTCATTTGGAGAGG |
| pOp6 | DEX Inducible promoter | AATTGTGAGCGCTCACAATTCTTTCTCTTCCCTTTCTTCTTTCTAGTCTTTCAATTGT GAGCGCTCACAATTCTTTCTCTTCCCTTTCTTCTTTCTAGTCTTTCAATTGTGAGCG CTCACAATTCTTTCTCTTCCCTTTCTTCTTTCTTTCAATTGTGAGCGCTCACAATTCT TTCTCTTCCCTTTCTTCTTTCTAGTCTTTCAATTGTGAGCGCTCACAATTCTTTCTCT TCCCTTTCTTCTTTCTAGTCTTTCAATTGTGAGCGCTCACAATTCTTTCTCTTCCCTT TCTTCTTTCTAGTGGATCGATCTTCGCAAGACCCTTCCTCTATATAAGGAAGTTCAT TTCATTTGGAGAGGA |
| NEV | Transcripti on Factor | ATGGCCCAGGCGGCCCTCGAGCCCGGGGAGAAGCCCTATGCTTGTCCGGAATGTG GTAAGTCCTTCAGCCAGAGCAGCAACCTGGTGCGCCACCAGCGTACCCACACGGG TGAAAAACCGTATAAATGCCCAGAGTGCGGCAAATCTTTTAGCCAGAGCAGCTCC CTGGTGCGCCATCAACGCACTCATACTGGCGAGAAGCCATACAAATGTCCAGAAT GTGGCAAGTCTTTCAGTCGCTCCGATAAACTGGTGCGCCACCAACGTACTCACACC GGTAAAAAAACTAGTGGCCAGGCCGGCCGCCGAAATGAAATGGGTGCTTCAGGA GACATGAGGGCTGCCAACCTTTGGCCAAGCCCTCTTGTGATTAAGCACACTAAGA AGAATAGCCCTGCCTTGTCCTTGACAGCTGACCAGATGGTCAGTGCCTTGTTGGAT GCTGAACCGCCCATGATCTATTCTGAATATGATCCTTCTAGACCCTTCAGTGAAGC CTCAATGATGGGCTTATTGACCAACCTAGCAGATAGGGAGCTGGTTCATATGATC AACTGGGCAAAGAGAGTGCCAGGCTTTGGGGACTTGAATCTCCATGATCAGGTCC ACCTTCTCGAGTGTGCCTGGCTGGAGATTCTGATGATTGGTCTCGTCTGGCGCTCC ATGGAACACCCGGGGAAGCTCCTGTTTGCTCCTAACTTGCTCCTGGACAGGAATC AAGGTAAATGTGTGGAAGGCATGGTGGAGATCTTTGACATGTTGCTTGCTACGTC AAGTCGGTTCCGCATGATGAACCTGCAGGGTGAAGAGTTTGTGTGCCTCAAATCC ATCATTTTGCTTAATTCCGGAGTGTACACGTTTCTGTCCAGCACCTTGAAGTCTCTG GAAGAGAAGGACCACATCCACCGTGTCCTGGACAAGATCACAGACACTTTGATCC ACCTGATGGCCAAAGCTGGCCTGACTCTGCAGCAGCAGCATCGCCGCCTAGCTCA GCTCCTTCTCATTCTTTCCCATATCCGGCACATGAGTAACAAAGGCATGGAGCATC TCTACAACATGAAATGCAAGAACGTTGTGCCCCTCTATGACCTGCTCCTGGAGATG TTGGATGCCCACCGCCTTCATGCCCCAGCCAGTCGCATGGGAGTGCCCCCAGAGG AGCCCAGCCAGACCCAGCTGGCCACCACCAGCTCCACTTCAGCACATTCCTTACA AACCTACTACATACCCCCGGAAGCAGAGGGCTTCCCCAACACGATCGGGCGCGCC GACGCGCTGGACGATTTCGATCTCGACATGCTGGGTTCTGATGCCCTCGATGACTT TGACCTGGATATGTTGGGAAGCGACGCATTGGATGACTTTGATCTGGACATGCTC GGCTCCGATGCTCTGGACGATTTCGATCTCGATATGTTAATTAACTACCCGTACGA CGTTCCGGACTACGCTTCTTGA |

| | | |
|---|---|---|
| LhGR | Transcription Factor | ATGGCTAGTGAAGCTCGAAAAACAAAGAAAAAAATCAAAGGGATTCAGCAAGCC<br>ACTGCAGGAGTCTCACAAGACACTTCGGAAAATCCTAACAAAACAATAGTTCCTG<br>CAGCATTACCACAGCTCACCCCTACCTTGGTGTCACTGCTGGAGGTGATTGAACCC<br>GAGGTGTTGTATGCAGGATATGATAGCTCTGTTCCAGATTCAGCATGGAGAATTAT<br>GACCACACTCAACATGTTAGGTGGGCGTCAAGTGATTGCAGCAGTGAAATGGGCA<br>AAGGCGATACCAGGCTTCAGAAACTTACACCTGGATGACCAAATGACCCTGCTAC<br>AGTACTCATGGATGTTTCTCATGGCATTTGCCCTGGGTTGGAGATCATACAGACAA<br>TCAAGTGGAAACCTGCTCTGCTTTGCTCCTGATCTGATTATTAATGAGCAGAGAAT<br>GTCTCTACCCTGCATGTATGACCAATGTAAACACATGCTGTTTGTCTCCTCTGAAT<br>TACAAAGATTGCAGGTATCCTATGAAGAGTATCTCTGTATGAAAACCTTACTGCTT<br>CTCTCCTCAGTTCCTAAGGAAGGTCTGAAGAGCCAAGAGTTATTTGATGAGATTCG<br>AATGACTTATATCAAAGAGCTAGGAAAAGCCATCGTCAAAAGGGAAGGGAACTC<br>CAGTCAGAACTGGCAACGGTTTTACCAACTGACAAAGCTTCTGGACTCCATGCAT<br>GAGGTGGTTGAGAATCTCCTTACCTACTGCTTCCAGACATTTTTGGATAAGACCAT<br>GAGTATTGAATTCCCAGAGATGTTAGCTGAAATCATCACTAATCAGATACCAAAA<br>TATTCAAATGGAAATATCAAAAAGCTTCTGTTTCATCAAAAATCTACTAGCAAACC<br>GGTAACGTTATACGACGTCGCTGAATACGCCGGCGTTTCTCATCAAACCGTTTCTA<br>GAGTGGTTAACCAGGCTTCACATGTTAGCGCTAAAACCCGGGAAAAAGTTGAAGC<br>TGCCATGGCTGAGCTCAACTACATCCCGAACCGTGTTGCGCAGCAGCTGGCTGGT<br>AAACAAAGCTTGCTGATCGGTGTCGCGACCTCGAGCTTGGCCCTGCACGCGCCGT<br>CGCAAATTGTCGCGGCGATTAAATCTCGCGCCGATCAACTGGGTGCCAGCGTGGT<br>GGTGTCGATGGTAGAACGAAGCGGCGTCGAAGCCTGTAAAGCGGCGGTGCACAA<br>TCTTCTCGCGCAACGCGTCAGTGGGCTGATCATTAACTATCCGCTGGATGACCAGG<br>ATGCCATTGCTGTGGAAGCTGCCTGCACTAATGTTCCGGCGTTATTTCTTGATGTC<br>TCTGACCAGACACCCATCAACAGTATTATTTTCTCCCATGAAGACGGTACGCGACT<br>GGGCGTGGAGCATCTGGTCGCATTGGGTCACCAGCAAATCGCGCTGTTAGCGGGC<br>CCATTAAGTTCTGTCTCGGCGCGTCTGCGTCTGGCTGGCTGGCATAAATATCTCAC<br>TCGCAATCAAATTCAGCCGATAGCGGAACGGGAAGGCGACTGGAGTGCCATGTCC<br>GGTTTTCAACAAACCATGCAAATGCTGAATGAGGGCATCGTTCCCACTGCGATGC<br>TGGTTGCCAACGATCAGATGGCGCTGGGCGCAATGCGCGCCATTACCGAGTCCGG<br>GCTGCGCGTTGGTGCGGATATCTCGGTAGTGGGATACGACGATACCGAAGACAGC<br>TCATGTTATATCCCGCCGTTAACCACCATCAAACAGGATTTTCGCCTGCTGGGGCA<br>AACCAGCGTGGACCGCTTGCTGCAACTCTCTCAGGGCCAGGCGGTGAAGGGCAAT<br>CAGCTGTTGCCCGTCTCACTGGTGAAAAGAAAAACCACTAGTGGATCGGAATTCG<br>CTAACTTCAACCAGTCCGGAAACATCGCTGATTCTTCCTTGAGCTTCACTTTCACT<br>AACTCTTCTAACGGACCTAACCTTATCACCACTCAGACCAACTCTCAGGCTCTTAG<br>CCAGCCAATCGCTAGCTCTAACGTGCACGACAACTTCATGAACAACGAGATCACT<br>GCTAGCAAGATCGATGATGGTAACAATTCTAAGCCTCTTAGCCCAGGATGGACTG<br>ATCAGACTGCTTACAACGCATTCGGTATCACTACCGGTATGTTCAACACCACTACC<br>ATGGACGATGTGTACAACTACCTCTTCGACGATGAGGATACTCCACCTAACCCTA<br>AGAAGGAGTGA |

**Table S2.** AD Primers used for TAIL PCR. W = A or T, S = G or C, N = A or T or G or C.

| Primer Name | Primer Sequence (5'-3') | Length | Degeneracy |
|---|---|---|---|
| AD1 | NGTCGASWGANAWGAA | 16 | 128 |
| AD2 | TGWGNAGSANCASAGA | 16 | 128 |
| AD3 | AGWGNAGWANCAWAGG | 16 | 128 |
| AD4 | STTGNTASTNCTNTGC | 16 | 256 |
| AD5 | NTCGASTWTSGWGTT | 15 | 64 |
| AD6 | WGTGNAGWANCANAGA | 16 | 256 |

**Table S3.** T-DNA left border Primer sequences for pGREENII0229 vector.

| Primer Name | Primer Sequence (5'-3') | Length |
|---|---|---|
| LB1 | TGGCATGACGTGGGTTTCTGGCAGCTGGACTTC | 33 |
| LB2 | ACAGGGCTTCAAGAGCGTGGTCGCTGTCATC | 31 |
| LB3 | TACATCGAGACAAGCACGGTCAACTTCCGTA | 31 |

**Table S4.** The p-values of two-sample two-sided t-test between fold changes of different treatments of Toggle 1.0 shoots (See Materials and Methods for plant numbers). Red color labels p-values larger than 0.05, thus not statistically significant differences. Mem is short for Memory. The difference in fold change between High Mem and High-Low is not statistically significant.

| | Low Mem | Low-High | Low-Low | Control | High Mem | High-High | High-Low |
|---|---|---|---|---|---|---|---|
| Low Mem | - | 0.0062 | 0.5565 | 0.0300 | 0.0004 | 0.0048 | 0.0007 |
| Low-High | - | - | 0.0020 | 0.0004 | 0.0001 | 0.0002 | 0.0001 |
| Low-Low | - | - | - | 0.0210 | 0.0001 | 0.0018 | 0.0004 |
| Control | - | - | - | - | 0.0010 | 0.0695 | 0.0022 |
| High Mem | - | - | - | - | - | 0.0062 | 0.7411 |
| High-High | - | - | - | - | - | - | 0.0109 |
| High-Low | - | - | - | - | - | - | - |

**Table S5.** The p-values of two-sample two-sided t-test between fold changes of different treatments of Toggle 1.0 roots (See Materials and Methods for plant numbers). Red color labels p-values larger than 0.05, thus not statistically significant differences. Mem is short for Memory. The difference in fold change between High Mem and High-Low is not statistically significant.

| | Low Mem | Low-High | Low-Low | Control | High Mem | High-High | High-Low |
|---|---|---|---|---|---|---|---|
| Low Mem | - | 0.0000 | 0.1598 | 0.0237 | 0.0000 | 0.0012 | 0.0000 |
| Low-High | - | - | 0.0000 | 0.0000 | 0.0000 | 0.0000 | 0.0000 |
| Low-Low | - | - | - | 0.0615 | 0.0000 | 0.0008 | 0.0000 |
| Control | - | - | - | - | 0.0000 | 0.0017 | 0.0000 |
| High Mem | - | - | - | - | - | 0.0001 | 0.2379 |
| High-High | - | - | - | - | - | - | 0.0000 |
| High-Low | - | - | - | - | - | - | - |

**Table S6.** The p-values of two-sample two-sided t-test between fold changes of different treatments of Toggle 2.0 shoots (See Materials and Methods for plant numbers). Red color labels p-values larger than 0.05, thus not statistically significant differences. Mem is short for Memory. The difference in fold change between High Mem and High-Low is not statistically significant.

| | Low Mem | Low-High | Low-Low | Control | High Mem | High-High | High-Low |
|---|---|---|---|---|---|---|---|
| Low Mem | - | 0.0000 | 0.0278 | 0.0470 | 0.0204 | 0.7834 | 0.0070 |
| Low-High | - | - | 0.0000 | 0.0000 | 0.0002 | 0.0001 | 0.0000 |
| Low-Low | - | - | - | 0.0037 | 0.0096 | 0.1386 | 0.0014 |
| Control | - | - | - | - | 0.0590 | 0.2208 | 0.0676 |
| High Mem | - | - | - | - | - | 0.0330 | 0.2349 |
| High-High | - | - | - | - | - | - | 0.0337 |
| High-Low | - | - | - | - | - | - | - |

**Table S7.** The p-values of two-sample two-sided t-test between fold changes of different treatments of Toggle 2.0 roots (See Materials and Methods for plant numbers). Red color labels p-values larger than 0.05, thus not statistically significant differences. Mem is short for Memory. The difference in fold change between the High Mem and High-Low is not statistically significant.

| | Low Mem | Low-High | Low-Low | Control | High Mem | High-High | High-Low |
|---|---|---|---|---|---|---|---|
| Low Mem | - | 0.0000 | 0.0643 | 0.3049 | 0.0000 | 0.9208 | 0.0000 |
| Low-High | - | - | 0.0000 | 0.0000 | 0.0000 | 0.0000 | 0.0000 |
| Low-Low | - | - | - | 0.0068 | 0.0000 | 0.4391 | 0.0000 |
| Control | - | - | - | - | 0.0000 | 0.6534 | 0.0000 |
| High Mem | - | - | - | - | - | 0.0001 | 0.1070 |
| High-High | - | - | - | - | - | - | 0.0001 |
| High-Low | - | - | - | - | - | - | - |

**Table S8.** The p-values of two-sample two-sided t-test between fold changes of different treatments of Toggle 2.1 shoots (See Materials and Methods for plant numbers). Red color labels p-values larger than 0.05, thus not statistically significant differences. Mem is short for Memory. The difference in fold change between the High Memory and High-Low is statistically significant.

| | Low Mem | Low-High | Low-Low | Control | High Mem | High-High | High-Low |
|---|---|---|---|---|---|---|---|
| Low Mem | - | 0.0000 | 0.8914 | 0.0777 | 0.0003 | 0.8806 | 0.0023 |
| Low-High | - | - | 0.0003 | 0.0000 | 0.0000 | 0.0001 | 0.0001 |
| Low-Low | - | - | - | 0.5699 | 0.0109 | 0.8462 | 0.0053 |
| Control | - | - | - | - | 0.0011 | 0.2137 | 0.0034 |
| High Mem | - | - | - | - | - | 0.0015 | 0.0356 |
| High-High | - | - | - | - | - | - | 0.0028 |
| High-Low | - | - | - | - | - | - | - |

**Table S9.** The p-values of two-sample two-sided t-test between fold changes of different treatments of Toggle 2.1 roots (See Materials and Methods for plant numbers). Red color labels p-values larger than 0.05, thus not statistically significant differences. Mem is short for Memory. The difference in fold change between the High Mem and High-Low is not statistically significant.

| | Low Mem | Low-High | Low-Low | Control | High Mem | High-High | High-Low |
|---|---|---|---|---|---|---|---|
| Low Mem | - | 0.0002 | 0.3041 | 0.3898 | 0.0002 | 0.0031 | 0.0002 |
| Low-High | - | - | 0.0003 | 0.0001 | 0.0000 | 0.0000 | 0.0000 |
| Low-Low | - | - | - | 0.5220 | 0.0005 | 0.0235 | 0.0004 |
| Control | - | - | - | - | 0.0001 | 0.0018 | 0.0002 |
| High Mem | - | - | - | - | - | 0.0010 | 0.1251 |
| High-High | - | - | - | - | - | - | 0.0007 |
| High-Low | - | - | - | - | - | - | - |

**Table S10.** Parameter values used for the choice of Toggle 1.0 and Toggle 2.0 components. These parameter values were estimated from the protoplast assays using Equation S3. Fold change indicates the ratio of the maximal expression level with no inducer to the lowest expression level achieved at the maximal inducer level. B, H and n are parameters characterizing the response functions of promoter-repressor pairs (defined in Equation S3). The parameter b is the dimensionless maximum expression level as defined in Equation S2, and 2b is the dimensionless maximum expression level of two copies of the LexA-based promoter to balance the stronger Gal4-based promoter. The choice of these combinations was based on the data from the best performing replicate, which are shown here.

| | **Construct Name** | **Fold Change** | **B ($10^8$)** | **H ($10^8$)** | **b** | **2b** | **n** |
|---|---|---|---|---|---|---|---|
| **Toggle 1.0** | $NOS_{2xGal4}$.EAR, Repressor 1 | 2.53 | 3.61 | 0.95 | 1.40 | N.A. | 3.03 |
| | $35S_{2xLexA}$.EAR, Repressor 2 | 5.99 | 0.66 | 2.57 | 0.69 | 1.38 | 2.56 |
| **Toggle 2.0** | $NOS_{2xGal4}$.EAR, Repressor 1 | 2.53 | 3.61 | 0.95 | 1.40 | N.A. | 3.03 |
| | $35S_{4xLexA}$TATA.OFPx, Repressor 2-OFPx | 4.31 | 0.48 | 1.01 | 0.51 | 1.02 | 3.31 |

**Table S11.** Parameter values estimated from the protoplast assays using Equation S4 with estimated Hill coefficient less than 6. Fold change indicates the ratio of the maximal expression level with no inducer to the lowest expression level achieved at the maximal inducer level. A, B, H and n are parameters characterizing the response functions of promoter-repressor pairs (Equation S4).

| **Constructs** | **Replicate Index** | **Fold Change** | **A ($10^8$)** | **B ($10^8$)** | **H ($10^8$)** | **N** |
|---|---|---|---|---|---|---|
| $NOS_{2xGal4}$.EAR | 3 | 2.53 | 1.704 | 3.264 | 1.279 | 2.615 |
| $NOS_{2xGal4}$.EAR | 4 | 2.04 | 0.7942 | 1.172 | 0.7805 | 1.783 |
| $35S_{2xLexA}$.EAR | 2 | 4.68 | 0.3856 | 1.914 | 1.203 | 3.406 |
| $35S_{2xLexA}$.EAR | 3 | 5.99 | 0.107 | 0.5607 | 1.543 | 3.678 |
| $35S_{4xLexA}$TATA.OFPX | 1 | 4.31 | 0.0733 | 0.3932 | 0.833 | 4.852 |

**SI Notes**

***1. Designing principles of a bi-stable toggle switch***

In order to model and simulate the plant toggle switch circuit, composed of two mutually repressing promoter-repressor pairs, we used a dimensionless system of ODEs as described [1]. The initial dimensional ODE equations can be written as follows. The rate of production of the repressor protein (X or Y) is determined by the concentration of its opposing repressor (Y or X) through a repressing Hill function, and its rate of degradation. Degradation is modeled as a first order process, yielding the following equations.

$$\frac{dX}{dT} = \frac{b_x}{1 + \left(\frac{Y}{K_x}\right)^{n_x}} - d_x X$$

$$\frac{dY}{dT} = \frac{b_y}{1 + \left(\frac{X}{K_y}\right)^{n_y}} - d_y Y$$

**Equation S1**

Here, X and Y are the concentrations of the two repressors that are mutually repressing each other in the toggle switch design; $b_x$ and $b_y$ are the maximum expression levels of the promoters producing X and Y respectively. $K_x$ and $K_y$ are the concentrations of X and Y that repress the promoter expression to its half-maximal level and $n_x$ and $n_y$ are Hill coefficients. The parameters $d_x$ and $d_y$ are the first-order degradation rate constants of X and Y, respectively.

We de-dimensionalized these two ODEs into the form shown in **Equation S2** to reduce the dimensionality of parameter space.

$$\frac{dx}{dt} = \frac{\tilde{b}_x}{1 + (y)^{n_x}} - x$$

$$\frac{dy}{dt} = \frac{\tilde{b}_y}{1 + (x)^{n_y}} - d_r y$$

**Equation S2**

Where:

$x = \frac{X}{K_y}$ and $y = \frac{Y}{K_x}$ are dimensionless concentrations of the two repressors X and Y;

$\tilde{b}_x = \frac{b_x}{K_y d_x}$ and $\tilde{b}_y = \frac{b_y}{K_x d_x}$ are the dimensionless maximum expression levels of the promoters producing X and Y respectively;

$n_x$ and $n_y$ are still the Hill coefficient of the promoters producing X and Y, respectively;

$d_r = \frac{d_y}{d_x}$ is the ratio between the degradation rate constant of X and Y, respectively. Time is de-dimensionalized by tdx.

We also assumed the degradation rate constants to be the same, thus $d_r = 1$. For simplicity, we relabel $\tilde{b}_x$ as $b_x$ and $\tilde{b}_y$ as $b_y$ in the remainder of the SI Notes. The resulting system of equations can be used to simulate and gain insights on the behaviors of toggle switch circuits. XPPaut (a tool to solve differential equations and perform bifurcation analysis https://sites.pitt.edu/~phase/bard/bardware/xpp/xpp.html), was used to generate the 2D phase diagrams of $b_x$ and $b_y$ at pre-determined values of Hill coefficients (Main text Figure 3).

From these 2D phase diagrams we can identify two principles for successful design of a toggle switch, which are the same as previously identified [1]. These are: 1) a sigmoidal repressor response with a high Hill coefficient, and 2) large and balanced values of the dimensionless maximum expression levels, as defined above in Equation S2.

These two principles can be understood by referring to Figure 3. As can be seen in the figure, if the value of the Hill coefficients, which measure the sigmoidal response, is larger, the bi-stable region is also larger. Secondly, larger and balanced values of the promoter strength, $b_x$ and $b_y$, are more likely to lie near the center of the bistable region, whereas smaller or unbalanced values are more likely, other things remaining constant, to lie outside, or near the edge of, the bistable region. Therefore, large and balanced $b_x$ and $b_y$ is another key principle for bistability.

### *2. Design of Toggle 1.0 and Toggle 2.0*

Building a toggle switch in plants requires genetic parts that produce bistable behavior when combined in the appropriate circuit topology. For the double repressor circuit topology [2], we can describe bistable bounds for the parameters of the transfer function of the two repressor-promoter pairs by two principles, as follows (previously identified (Gardner, 2000 #1770), and described here in more detail). A successful bistable toggle switch requires 1) a sigmoidal repressor response with a high Hill coefficient, and 2) High expression levels, from each promoter, that are balanced and mathematically made dimensionless. The latter is explicitly defined below in Equation S3 (below)

We have previously published our methods for construction and characterization of a library of repressor-promoter pairs [3]. Comparison of quantitative parameters estimated from transient expression of components in protoplasts with those estimated from stable expression in plants yielded parameters that were approximately within a factor of two of each other, with the discrepancies most likely due to effects arising from the random integration of the synthetic circuit in the plant genome. This level of possible discrepancies led us to conclude that while quantitative predictions based on these parameters were likely to be subject to significant error, we could still use the two bistability criteria defined above to design a mutual repression-based toggle switch. The first putative combination to satisfy the required criteria very early on was the combination consisting of $NOS_{2xGal4}$.EAR and $35S_{2xLexA}$.EAR, which had Hill coefficients above 3 and 2, respectively (Table S10). However, the Gal4-based repressible promoter was found to have a higher maximal expression, and to balance the promoter strengths we combined two copies of the LexA-based promoter with one copy of the Gal4-based promoter (Table S10). This construct is described as Toggle 1.0 in the text. Given the time requirements for production of plants with stable integration of synthetic genetic circuits, we began to assemble this construct in plants while we continued testing many other promoter-repressor pairs. After testing about 120 such genetic circuits, the high sigmoidality criteria led to identification of three promoter-repressor pairs where at least one experimental replicate was fit by a Hill function with a Hill coefficient, n, greater than

3 ($35S_{4xLexA}$TATA.OFPx, $_{2xGal4}$NOS.EAR and $NOS_{2xGal4}$.EAR). Of these three, the combination consisting of two copies of $35S_{4xLexA}$TATA.OFPx and one copy of $NOS_{2xGal4}$.EAR paired the best in terms of balanced dimensionless maximum expression levels, since here too the Gal4-based promoter was found to be stronger (Table S10). This genetic circuit was assembled and is labeled as Toggle 2.0 in the text.

#### *3. Data Analysis of Toggle 1.0*

Luminescence heat maps of whole plants carrying Toggle 1.0 are shown in Figure S4**.** The plots were reconstructed using the raw luciferase luminescence images and the ROIs acquired from the image processing as described in the *SI Materials and Methods*. One plant in each treatment is shown as an example, whereas data are collected from a minimum of three genetically identical plants per treatment. From analysis of the data such as that shown in the heat maps, the Low and High states were well established and the switching behaviors between the two steady states were as expected. However, the High Memory state did not show stability, with no obvious differences from the High to Low treatment. Interestingly, the roots seemed to perform better than shoots in terms of luminescence intensity and switching behavior.

These observations were supported by quantitative data showing the temporal luminescence intensity profiles (Figure S5a & b) and population mean fold change (Figure S5c & d). Figure S5a & b shows the intensity profile of shoots and roots, respectively, as extracted from the heatmaps (Figure S4). The behavior of the toggle switch showed some significant quantitative differences between shoots and roots. The mean fold change of Low to High was 21.3-fold for roots compared to 2.3-fold for shoots. For High to Low, the mean fold change of roots was 24.9-fold, and 3.9-fold for shoots. However, both shoots and roots failed to establish a stable High State as predicted for a toggle switch, as the differences between the High Memory and High to Low are not statistically significant from the 2-sample t-test.

#### *4. MCMC Parameter Estimation Results for Toggle 1.0*

Shoots: (i) Fits to the means: 239 parameter sets met the criteria for good parameter sets (discussed in the *SI Materials and Methods*), of which 2 were bistable and an additional 3 were indistinguishable from bistable. Thus, about 2.1% of parameter sets showed bistability for the duration of the experiment. Of these 239 parameter sets, 90 were classified as best parameter sets and they included all 5 parameter sets that showed bistability for the duration of the experiment (5.56%).
(ii) Pooled fits to each technical replicate: 167 parameter sets met the criteria for good parameter sets, of which 6 were bistable and 2 were indistinguishable from bistable (for a total of 4.8%). When using this random sampling of replicates method, we only report results of all the good parameter sets.
(iii) Fits to the data of Figure S4: We obtained 141 good parameter sets of which 0 were bistable.

Roots: (i) Fits to the means: We found 325 good parameter sets of which 0 were bistable.
(ii) Pooled fits to each technical replicate: Of 100 good parameter sets, 3 were bistable and none were indistinguishable from bistable.

(iii) Fits to the data of Figure S4: We found 244 good parameter sets of which 0 were bistable.

Based on these results we classified Toggle 1.0 as a monostable system.

***5. Data Analysis of Toggle 2.0***

The intensity plots shown in Figure S7c & f, shows that the Low and High states of Toggle 2.0 were well established. The High state of shoots was maintained even after the 4-OHT inducer was removed, as expected for a functional toggle switch (Figure S7e). However, the addition of DEX inducer to switch from the High to the Low state (Figure S7g) did not produce an effective switch, and in fact the luminescence on the final day was not statistically significantly different from the High Memory (Figure S7e). Interestingly, and distinct from Toggle 1.0, the shoots had some performance which was better in Toggle 2.0 than in Toggle 1.0, with a mean fold change of 37.5 (compared to a fold-change of 2.3 in Toggle 1.0), as shown in Figure S8c. On the other hand, the roots of Toggle 2.0 performed similarly to Toggle 1.0. It is noteworthy that the Low to High change in luminescence was almost two orders of magnitude, a fold-change of 80 (Figure S8d).

***6. MCMC Parameter Estimation Results for Toggle 2.0***

Shoots: (i) Fits to the means: We obtained 1,141 good parameter sets (after running the MCMC algorithm 10,000 times), of which 278 were bistable and 822 were indistinguishable from bistable (totaling 96.4%). Out of these, 1,079 were best parameter sets, of which 260 were bistable and 812 indistinguishable from bistable (totaling 99.4%).
(ii) Pooled fits to each technical replicate: Of 111 good parameter sets, 22 were bistable and 84 were indistinguishable from bistable, totaling 95.5%.
(iii) Fits to data of Figure S7: Almost all the good parameter sets here were also best (142/146). Of the 146 good sets, 21 were bistable and 121 were indistinguishable from bistable (in total 97%).

Roots: (i) Fits to the means: 134 parameter sets were good (and best); of these none were bistable.
(ii) Pooled fits to each technical replicate: 134 parameter sets were good (and best), and none of these was bistable.
(iii) Fits to the data of Figure S7: Almost all the parameter sets were best (129/132). Out of 132 good parameter sets, 1 was bistable and 1 indistinguishable from bistable.

The MCMC fits therefore suggest that the Toggle 2.0 circuit is behaving like a bistable toggle switch in the shoots and a monostable system in the roots. However, since the luminescence change from High to Low in the shoots was not statistically significant, we surmised that the inducible promoter driving expression of the GEAR repressor is not sufficiently strong (our data do not distinguish whether it is the promoter or the GEAR repressor). Toggle 2.1 therefore was constructed with two copies of this inducible promoter driving expression of the GEAR repressor as detailed in *SI Materials and Methods*.

***7. Data Analysis of Toggle 2.1***

The circuit design with Toggle 2.0 along with a second copy of the DEX-inducible Gal4EAR repressor is referred to as Toggle 2.1. From the intensity plots shown in Figure S11, the Low and

High states are again clearly distinguishable in Toggle 2.1. In accordance with the modeling predictions, the stability of the High state was maintained, and the switching behavior from High to Low was improved in shoots with a fold change of 6.4 compared to 2.6 in Toggle 2.0 (Figure S11c). This difference is statistically significant (Table S8).

The roots still performed similarly to Toggle 2.0 (Figure S11a & b). The fold change of Low to High was 18.9 and the fold change of High to Low was 10.7 (Figure S11d).

#### *8. MCMC Parameter Estimation Results for Toggle 2.1*

Shoots: (i) Fits to the means: Out of 321 good parameter sets, 113 were bistable and 206 were indistinguishable from bistable (totaling 99.4%). Of the good fits, 70 parameter sets met the definition of best parameter sets, and 58 of these were bistable and 12 indistinguishable from bistable (totaling 100%).
(ii) Pooled fits to each technical replicate: Out of 148 good parameter sets, 54 were bistable and 85 indistinguishable from bistable (totaling 94%).
(iii) Fits to the data in Figure S10: Out of 150 good parameter sets, 68 were bistable and 82 indistinguishable from bistable (totaling 100%). Of these 150, we found 71 best parameter fits, of which 41 were bistable and 30 were indistinguishable from bistable (totaling 100%).

Roots: (i) Fits to the means: We found 133 good parameter sets out of which 17 were bistable and 4 were indistinguishable from bistable (totaling 15.8%). Of these 133, 86 were best parameter sets and they included the 21 parameters that were bistable for the duration of the experiment (24%).
(ii) Pooled fits to each technical replicate: We ran the MCMC procedure 1,700 times and found 354 good parameter sets. Out of these 87 were bistable and 25 indistinguishable from bistable (totaling 31.6%). (ii) Fits to the data in Figure S10: 155 out of 310 good parameters were bistable and 11 were indistinguishable from bistable (totaling 53.6%). However, 201 of these 310 were best parameters, and of these, 152 were bistable and 10 were indistinguishable from bistable, or a total of 80.6% were bistable for the duration of the experiment.

The data strongly suggest that shoots containing Toggle 2.1 have bistability. The behavior of the circuit in the roots is more ambiguous (see main text for discussion).

#### *9. Transgene Insertion and Copy Number Estimation*

Thermal asymmetric interlaced polymerase chain reaction (TAIL-PCR) has been used to retrieve sequences flanking insertion sites and estimate the number of insertions in plant chromosomes [4]. Here, we followed standard TAIL-PCR protocol as reported in Singer and Burke [4]. In brief, three subsequent rounds of TAIL-PCR were performed using arbitrary degenerate (AD) primers (Table S2) and nested insertion-specific primers (Table S3). We used published arbitrary degenerate primers from *Arabidopsis thaliana* [4]. The nested insertion-specific primers were designed targeting the BAR gene (selection marker) in the pGREENII0229 binary vector. Both shoot and root genomic DNAs extracted from Toggle 2.0 homozygous lines were used as a template. Only secondary and tertiary TAIL-PCR products were analyzed on gel. The result showed a single band shifting between secondary and tertiary products in shoot as well as root DNA, suggesting a single copy of the transgene (Figure S12a). Subsequently, the gel-purified tertiary TAIL-PCR products were cloned in the pJET vector and sequenced. The results show that all the sequenced products contained the inserted T-DNA and its flanking genomic sequences of the Arabidopsis genome (Figure S12b). The transgene inserted in Chromosome 5, in the intergenic region between AT5G39980 and AT5G39990. Generation of clean DNA sequence products from TAIL PCR off the insert's left border is consistent with one intact T-DNA in these plants.

***10. Comparison of parameters estimated from protoplast assays and whole plant luminescence data***

In a previously published work, we have developed a method to quantitatively characterize repressible promoters using high throughput protoplast assays[1]. We developed a statistical model to remove intrinsic experimental variability, and quantitative promoter values were characterized for over 120 promoter-repressor pairs in *Arabidopsis* and over 100 pairs in sorghum. For the purpose of this study, the promoter-repressor pairs in *Arabidopsis* are of interest. The performance of the repressible promoter was quantified by fitting the data to a Hill function as shown in Equation S3.

$$L_i = \frac{B_i}{1 + \left(\frac{F_i}{H_i}\right)^{n_i}} \qquad \textbf{Equation S3}$$

Here $L_i$ is the output of the repressible promoter, measured in units of molecular concentration, and $F_i$ is the molecular concentration of the repressor protein. The subscript i indicates the $i^{th}$ promoter-repressor in the library. Molecular concentration of $L_i$ were calculated from luciferase luminescence values, converted into concentrations by using a calibrated standard curve between concentration and luminescence [3]. As typical for repressing Hill functions [5] the parameter $B_i$ represents the maximum promoter strength, the parameter $H_i$ is the repressor concentration at half-maximal expression and the parameter $n_i$ (Hill coefficient) represents the sigmoidality of the response curve.

The raw data generated by protoplast assays were processed and normalized following the procedures described [3]. The normalized data were thereafter fitted to Equation S3 using a nonlinear least-squares package in Matlab (Mathworks).

As detailed in the *Materials and Methods*, a simple ODE system (Equation 1) was developed to describe the experimental results of the stably transformed toggle switch constructs and quantitative parameter values can be estimated by maximizing a log-likelihood using a MCMC scheme. These parameter values were then fed into Equation S3 without the inducible terms to evaluate their bistability. Such parameterized models can also be used to test out different strategies to modify the design of Toggle 2.0. In addition, it is also possible to compare these parameter values to the ones previously estimated from the protoplast assays characterizing the repressor-promoter combinations. Such comparisons will help us to evaluate the accuracy of the transient expression in the protoplast assays as an approximation to the stable transformations.

To make the parameter values estimated from the two sources comparable, the method described above in parameter estimation section needs some modifications. First, one term characterizing basal expression was added into Equation S3.

$$L_i = A_i + \frac{B_i}{1 + \left(\frac{F_i}{H_i}\right)^{n_i}} \qquad \textbf{Equation S4}$$

where $A_i$ represents the relative basal expression of the $i^{th}$ promoter and all the other variables are defined as in parameter estimation section. Each of these parameters quantitatively characterizes how the promoter repressor pair behaves.

The raw data generated by protoplast assays were processed and normalized following the same procedures described [3]. The normalized data were then fitted to Equation S4 using a nonlinear least-squares package in Matlab (Mathworks). Specifically, a Robust option of Least Absolute Residuals (LAR) and the Trust-Region algorithm were used. Parameter values were constrained to positive values for fitting. A good fit was defined to have an estimated value of Hill coefficient no larger than 6, which is assumed to be biologically relevant in this study. Out of the replicates of the $NOS_{2xGal4}$.EAR, $35S_{2xLexA}$.EAR and $35S_{4xLexA}$TATA.OFPx, the estimated parameter values of good fits are listed in **Table S11**.

To directly compare the parameter values estimated from the protoplast assays with the whole plant luminescence data, we used de-dimensionalized parameter values by normalizing A and B by their corresponding H of the same promoter-repressor combination. For a brief proof that these are equivalent and comparable, please refer to the next section. As shown in Figure S13, the parameter values estimated from the protoplast assays generally overlap with the distribution of dimensionless parameter values estimated from the whole plant luminescence data. While this result supports the use of protoplasts for quantitative characterization of plant plants for synthetic biology, the possible quantitative differences between the protoplast estimates and those in a plant can be quite high, as seen in the figure. Therefore, quantitative predictions, for example of bistability properties, based on protoplast data are still subject to significant error. Allowing for these margins of error and using protoplast data for qualitative and semi-quantitative predictions however, allowed us to overcome these limitations and construct the first genetic toggle switch in plants.

### *12. Proof of the comparability of de-dimensionalized parameter values estimated from protoplast assays and whole plant luminescence data*

In our previous work [3], we developed a quantitative model of the parameters estimated from the protoplast assays in terms of the response function at the single-plasmid level, and mathematical expressions connecting the single-plasmid level and the experimental observables. Using variables as defined in Equation S3, the fitting results can be represented as:

$$B_i = C_2 \langle N_{ij} \rangle_j \alpha_i \beta_i$$

$$H_i = C_1 \tilde{C} \langle N_{ij} \rangle_j \alpha_i K_i$$

The meaning of each parameter is described briefly here for the reader's convenience. Please refer to (1) for more details.
$\beta_i$ represents the maximal expression of the R-luciferase (Rluc) protein with no repressor in a single plasmid (i.e. promoter strength);
$K_i$ is the repressor concentration required for half-maximal expression of Rluc in a single plasmid (i.e. repressibility);
$\alpha_i$ represents the batch variability factor;
$N_{ij}$ represents the total number of plasmids in the j-th well of the i-th plasmid;
Therefore, $\alpha_i N_{ij}$ describes the plasmid copy number in viable protoplasts in the j-th well of the i-th plasmid.
$C_1$ is the concentration - luminescence proportionality factor of F-Luciferase (1);
$C_2$ is the concentration - luminescence proportionality factor of RLuc;
$\tilde{C}$ is a proportionality factor between the concentrations of repressor and FLuc (both of which are controlled by the same promoter).

A similar representation can be derived for the basal expression level A due to its similarity to $B_i$.

$$A_i = C_2 \langle N_{ij} \rangle_j \alpha_i a_i$$

Note that $a_i$ is the basal expression level in single-plasmid level, while $\alpha_i$ (alpha) is the batch variability factor. The mathematical representations of these parameters indicate an intuitive method to de-dimensionalize these parameters by simply dividing $A_i$ and $B_i$ by $H_i$.

$$\frac{A_i}{H_i} = \frac{C_2 a_i}{C_1 \tilde{C} K_i}$$

**Equation S5**

$$\frac{B_i}{H_i} = \frac{C_2 \beta_i}{C_1 \tilde{C} K_i}$$

As described [3], $C_1$ and $C_2$ can be estimated from the experimental standard curve relating the luminescence levels and the molecular numbers of the two types of luciferase. $C_1$ and $C_2$ were first estimated from standard curves measured experimentally. Then, their values were used to convert the experimental data from luminescence level to molecular numbers, after which the nonlinear least square fitting was carried out. Therefore, $A_i$, $B_i$ and $H_i$ are already in the unit of molecular numbers and the $C_1$ and $C_2$ in the dimensionless representation can be discarded. We can reasonably assume $\tilde{C} = 1$, meaning a 1:1 ratio between the molecular numbers of FLuc and repressor, because they are driven by the same promoter. Therefore, we have:

$$\frac{A_i}{H_i} = \frac{a_i}{K_i}$$

$$\frac{B_i}{H_i} = \frac{\beta_i}{K_i}$$

Following a similar logic, we can also develop a quantitative model of the parameters estimated from the whole plant luminescence data. Firstly, the mean luminescence emitted from a single pixel can be represented as:

$$\langle F_i \rangle = C_1 \tilde{C} \langle N_i \rangle R_i$$

where the meaning of each variable is similar to the protoplast assay but modified accordingly.
$F_i$ represents the luminescence emitted from a single pixel and the square brackets indicate population mean across all pixels included to quantify the luminescence.
$R_i$ represents the total number of repressors in a single cell;
$N_i$ represents the total number of plant cells in one pixel;
$C_1$ is the concentration - luminescence proportionality factor of FLuc;
$\tilde{C}$ is a proportionality factor between the repressor concentration and FLuc concentration (both of which are controlled by the same promoter), which we assume equals 1, as explained above.

Because FLuc was the only reporter of the toggle switch assembled and tested in the whole plant, and population mean luminescence per pixel was used for the nonlinear least-square fitting, the ODE system formulated in Equation 1 can be rescaled and the fitted parameters can be represented by:

$$A_i = C_1 \tilde{C} \langle N_i \rangle a_i$$

$$B_i = C_1 \tilde{C} \langle N_i \rangle \beta_i$$

$$H_i = C_1 \tilde{C} \langle N_i \rangle K_i$$

We can now de-dimensionalize the parameters in the same way.

$$\frac{A_i}{H_i} = \frac{a_i}{K_i}$$

$$\frac{B_i}{H_i} = \frac{\beta_i}{K_i}$$

Therefore, the de-dimensionalized parameter values estimated from the protoplast assay and the whole plant luminescence data are directly comparable.

#### *13. Testing the MCMC Algorithm Using Simulated Experimental Data*

We tested the reliability of the MCMC algorithm using a simulated dataset. We first picked a parameter set of the ODE model representing a bistable toggle switch, then simulated different treatments as applied in the experiments and sampled several discrete time points with a constant time step from the numerical solutions as in the experimental setup. Only one side of the toggle switch was sampled and used for the MCMC, as only one side of the toggle switch is labeled with the luciferase gene reporter in the experiments.

### SI References


[1] Gardner, T. S., Cantor, C. R., and Collins, J. J. (2000) Construction of a genetic toggle switch in Escherichia coli, *Nature 403*, 339-342.

[2] Lebar, T., Bezeljak, U., Golob, A., Jerala, M., Kadunc, L., Pirš, B., Stražar, M., Vučko, D., Zupančič, U., Benčina, M., Forstnerič, V., Gaber, R., Lonzarić, J., Majerle, A., Oblak, A., Smole, A., and Jerala, R. (2014) A bistable genetic switch based on designable DNA-binding domains, *Nature Communications 5*, 5007.

[3] Schaumberg, K. A., Antunes, M. S., Kassaw, T. K., Xu, W., Zalewski, C. S., Medford, J. I., and Prasad, A. (2016) Quantitative characterization of genetic parts and circuits for plant synthetic biology, *Nature Methods 13*, 94-100.

[4] Singer, T., and Burke, E. (2003) High-throughput TAIL-PCR as a tool to identify DNA flanking insertions, *Methods Mol Biol 236*, 241-272.

[5] Alon, U. (2006) *An Introduction to Systems Biology: Design Principles of Biological Circuits*, Taylor & Francis.
